# A novel metalloprotease family discovered through a tardigrade

**DOI:** 10.64898/2026.10.27.762089

**Authors:** Tomoka Higashi, Yohta Fukuda, Tsuyoshi Inoue

## Abstract

Tardigrades possess numerous proteins with poorly understood functions, including candidates proposed to have been acquired through horizontal gene transfer. Here, we characterized RvY_14254 from *Ramazzottius varieornatus*, previously designated ATPM5, whose homologs are widely distributed across bacteria and eukaryotes but remain largely uncharacterized. We determined the crystal structure of the protein, hereafter termed *Rv*Peti, at 1.35 Å resolution. *Rv*Peti adopts a compact, C-shaped architecture with no close structural homologs in the Protein Data Bank and contains a conserved His-Asp-Asp (HDD) metal-binding motif within a deep surface pocket. Structural comparison revealed partial similarity of the active-site architecture to that of metalloproteases, including pseudoalterin. Anomalous diffraction analyses demonstrated that the HDD site can accommodate Zn² and Mn² . Biochemical assays showed that *Rv*Peti exhibits Zn² -dependent proteolytic activity, which was strongly reduced by a metal chelating reagent and by alanine substitutions of the metal-coordinating residues. In contrast, only weak esterase activity was detected, with the substrates tested. These results identify *Rv*Peti as a founding member of a previously uncharacterized family of metalloproteases and provide a structural framework for investigating the physiological and evolutionary roles of this widespread protein family.

**statement:** Tardigrades, organisms capable of surviving extreme environment stress, possess numerous poorly characterized proteins. We determined the crystal structure of one such protein and found that it adopts a novel fold containing a metal-binding site. Biochemical analyses confirmed zinc-dependent proteolytic activity that was lost upon mutation of the metal-binding residues. These findings define a previously unrecognized metalloprotease family conserved across bacteria and eukaryotes, providing a structural framework for future function studies.

## 1. Introduction

Tardigrades, also known as water bears, are small but complex multicellular animals, typically less than 1 mm in body length. They inhabit soil, water, and the surfaces of plants, and can even be found living in the moss growing along roadsides (Nelson, 2002). Some tardigrades are known for a remarkable ability: under desiccating conditions, they reduce their body water content to only a few percent and form a compact, tun-shaped structure, entering a specific state known as anhydrobiosis (Wełnicz et al., 2011; Guidetti et al., 2011). In this extremely dehydrated state, tardigrades cease essentially all metabolic activity, appearing as if dead. However, the addition of water allows them to rapidly revive and resume normal activity. Remarkably, in the anhydrobiotic state, tardigrades can withstand severe environmental stresses such as desiccation (Rebecchi et al., 2007), extreme temperatures (Becquerel, 1950; Hengherr et al., 2009), high pressure (Ono et al., 2008; Seki & Toyoshima, 1998), and radiation (Horikawa et al., 2013, 2006). The molecular mechanisms underlying tolerance systems of tardigrades have long been studied for their potential to inform our understanding of survival strategies in life more broadly (Møbjerg & Neves, 2021). Moreover, these findings are expected to have applications in medicine and biotechnology, including the stabilization of biomolecules and the long-term preservation of cells and other biological materials (Kasianchuk et al., 2023; Lim et al., 2024).

To elucidate the molecular basis of this tolerance, transcriptomic and other omics approaches have been applied to multiple tardigrade species (Kamilari et al., 2026). Among many kinds of tardigrades, *Ramazzottius varieornatus*, a tardigrade capable of tolerating rapid desiccation, has been shown by transcriptome analysis to express numerous genes specifically in either the active or anhydrobiotic state (Hashimoto et al., 2016; Yoshida et al., 2017). Several tardigrade-specific proteins implicated in environmental stress tolerance have been characterized biochemically and structurally, including cytoplasmic-abundant heat-soluble (CAHS), secretory-abundant heat-soluble (SAHS), and damage suppressor (Dsup) proteins (Yamaguchi et al., 2012; Hashimoto et al., 2016; Fukuda et al., 2017; Boothby et al., 2017). Nevertheless, much remains to be understood about the molecular repertoire underlying the remarkable stress tolerance of tardigrades. One question that has attracted considerable attention is the contribution of proteins acquired through horizontal gene transfer (HGT). An initial analysis of the draft genome of *Hypsibius dujardini* (subsequently reidentified as *Hypsibius exemplaris*) suggested an unusually high frequency of HGT, with approximately one-sixth of its genes proposed to be of non-metazoan origin (Boothby et al., 2015). Subsequent analyses using improved genome assemblies and more stringent approaches to exclude contamination substantially revised this estimate downward and found no evidence for exceptionally extensive HGT in tardigrades (Koutsovoulos et al., 2016; Hashimoto et al., 2016). However, the rejection of extensive genome-wide HGT does not exclude the possibility that individual horizontally acquired genes have played important roles in tardigrade adaptation. Indeed, recent studies have identified putatively horizontally acquired proteins with potential roles in stress responses, including a radiation-responsive DOPA dioxygenase that promotes radiotolerance through betalain production (Li et al., 2024) and tardigrade ferritin-like metabolite phosphatases implicated in stress adaptation (Kato et al., 2024). Thus, although HGT is unlikely to represent a pervasive genomic basis for tardigrade extremotolerance, the extent to which specific HGT-derived proteins have contributed to the evolution of extreme stress tolerance remains an important open question. Despite the growing number of genes proposed to have originated through HGT, only a limited fraction of their protein products have been characterized experimentally, and their biochemical functions, structural properties, and physiological roles remain largely unexplored.

Here, we focused on RvY_14254 (GenBank ID: BAQ592.1; UniProt ID: A0A0E4AVP5), a candidate HGT-derived gene from *R. varieornatus* that is highly expressed in both the active and anhydrobiotic states (Hashimoto et al., 2016). The protein product of this gene is relatively small, with a molecular weight of only 14 kDa. Homology searches based on its amino acid sequence indicate that it is conserved across a wide range of species from eukaryotes to bacteria (Fig. 1A), suggesting that it plays some functional role in those organisms. Nevertheless, the structure, function, and physiological roles of this protein family remain unknown. RvY_14254 was previously predicted to localize to mitochondria and hence has been referred to as tardigrade-unique protein with predicted mitochondria targeting peptide 5 (ATPM5) (Tanaka et al., 2015). However, Tanaka *et al*. reported that the protein expressed in human HEK293T cells showed no mitochondrial localization. Its true localization and function were never clarified, and no further analysis has been conducted so far.

**Fig. 1.**
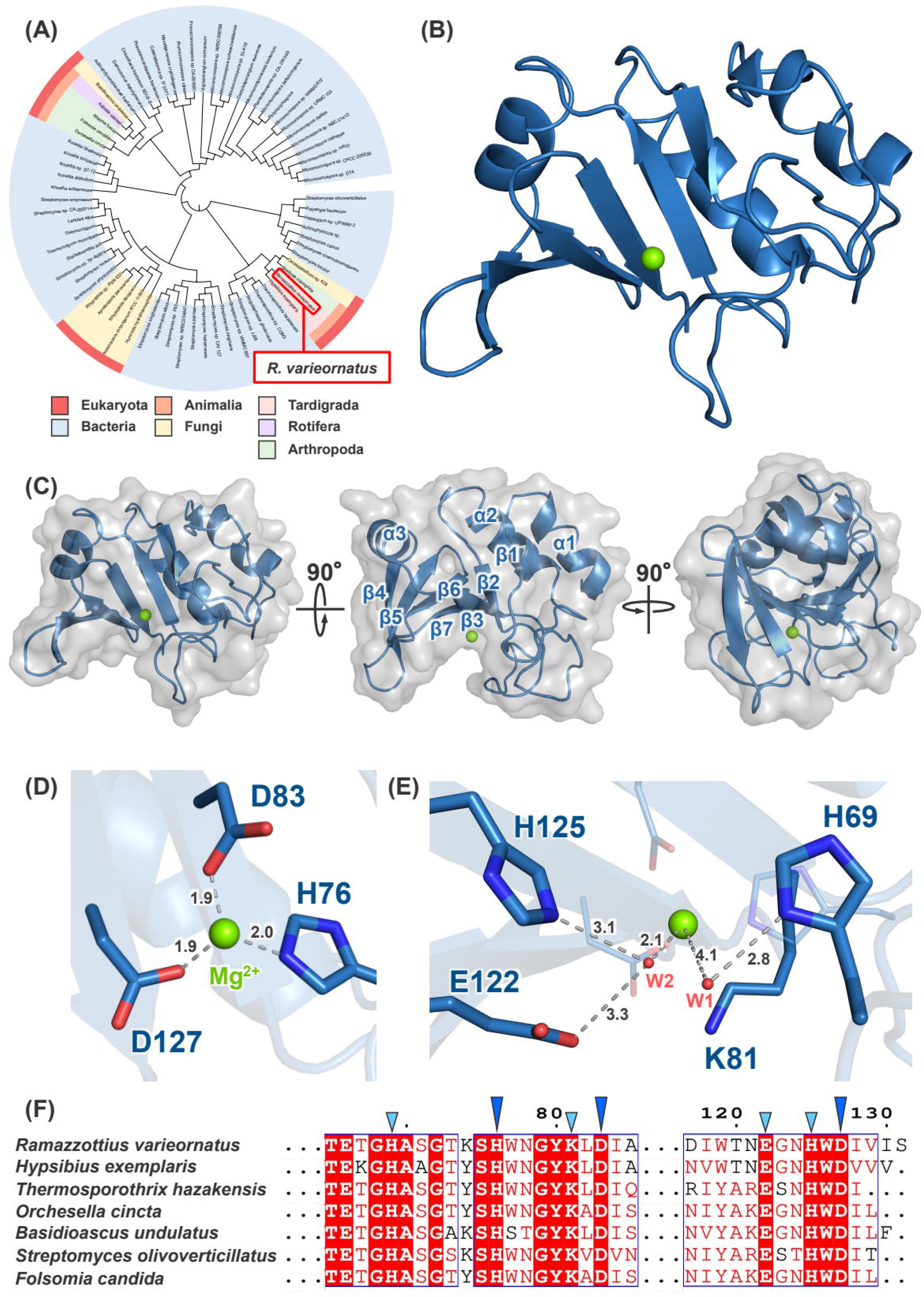
Overall structure of *Rv*Peti and its conserved active site. (A) Phylogenetic tree of *Rv*Peti. (B) Cartoon representation of the overall structure of *Rv*Peti. *Rv*Peti and Mg² ion are colored in blue and light green, respectively. (C) Surface representation of *Rv*Peti (gray) shown in three orientations, each rotated 90° from the previous view. α-helices and β-strands are numbered sequentially from the N-terminus. (D) Close-up view of the HDD motif. His76, Asp83, and Asp127 are shown as sticks. (E) Residues in the vicinity of the HDD motif. His69, Lys81, Glu122, and His125 are shown as sticks. (F) Multiple sequence alignment of *Rv*Peti and its top six homologs identified by BLAST. Blue and light blue triangles indicate the HDD motif residues and their neighboring residues, as shown in (D) and (E), respectively.

In this study, we determined the crystal structure of the RvY_14254 protein product and investigated its enzymatic properties. Our findings reveal a previously uncharacterized structure and provide insights into the metal-dependent enzymatic properties of this protein. We call the protein product of RvY_14254 *Rv*Peti (ProtEase from TardIgrade from *R. varieornatus*).

## 2. Results

### 2.1. Crystal structure of *Rv*Peti

We performed X-ray crystallographic analysis of *Rv*Peti. The crystal structure of the as-purified form was determined at 1.35 Å resolution (Table. 1, Fig. 1B). *Rv*Peti adopts a C-shaped overall architecture composed of seven β-strands and three α-helices, which together form a large groove-shaped open pocket at the center of the molecule (Fig. 1C).

Structural similarity searches against the Protein Data Bank (PDB) database using Foldseek (van Kempen et al., 2024) and the DALI server (Holm, 2022) revealed no close structural homologs of *Rv*Peti. Foldseek identified no protein structures with a TM-score ≥ 0.5, and the maximum Z-score from DALI was 7.0, reflecting only partial similarity at the level of local core structures. These results suggest that *Rv*Peti adopts a distinct overall fold, despite local structural similarities to known proteins.

Within the pocket formed by the β-sheets, we identified a metal-binding site formed by a His76-Asp83-Asp127 (HDD) motif (Fig. 1D). The electron density indicated the presence of a light rather than a heavy metal atom, and the site was consequently modeled as a putative Mg^2+^ site. The residues of the HDD motif may form part of an active site responsible for the metal-dependent enzymatic activity of *Rv*Peti. Near the HDD motif, polar and charged residues (His69, Lys81, Glu122, and His125) are solvent-exposed within the cleft. Two water molecules, Water 1 (W1) and Water 2 (W2), were also identified (Fig. 1E). W1 coordinates to the metal ion and forms a hydrogen bond network with His69, although W1 is farther from the metal ion (∼ 4.1 Å). W2 forms a hydrogen bond network with Glu122 and His125. The distance between W1 and W2 is about 2.7 Å. Most of the residues that constitute the active site are highly conserved across species, strongly suggesting their importance (Fig. 1F). If *Rv*Peti is an enzyme, these residues and water molecules may play roles in substrate recognition and stabilization and/or potentially serve as a proton transfer pathway during catalysis.

### 2.2. Partial structural similarity to a metalloprotease and functional prediction

To elucidate the function of *Rv*Peti, we manually searched the PDB for known protein structures similar to *Rv*Peti. This search identified pseudoalterin (PDB: 6IK4) from the deep-sea bacterium *Pseudoalteromonas* sp. CF6-2. Pseudoalterin is an M23 metalloprotease with a His-His-Asp (HHD) motif at its catalytic center. This protease was originally reported as an elastin-degrading enzyme exhibiting high hydrolytic activity toward bovine elastin (Zhao et al., 2012) and it was later shown that this enzyme also cleaves peptide bonds within peptidoglycan (Tang et al., 2020). Interestingly, a BLAST search based on *Rv*Peti retrieves a protein from a thermophilic bacterium *Thermosporothrix hazakensis* that is predicted to possess a putative peptidoglycan-binding domain.

*Rv*Peti and pseudoalterin share a similarly large, open-mouthed pocket forming the active site, as well as a comparable β-sheet arrangement (Figs. 2A, 2B, 2C). In contrast, their amino acid sequences and overall topology differ substantially (Fig. 2D), with *Rv*Peti sharing only 23.3% sequence similarity with pseudoalterin. *Rv*Peti possesses a longer loop forming the pocket, resulting in a more deeply buried active site. Furthermore, although the metal coordination distances are comparable, *Rv*Peti possesses an HDD motif whereas pseudoalterin possesses an HHD motif, reflecting a difference in one His/Asp residue (Fig. 2B).

**Fig. 2.**
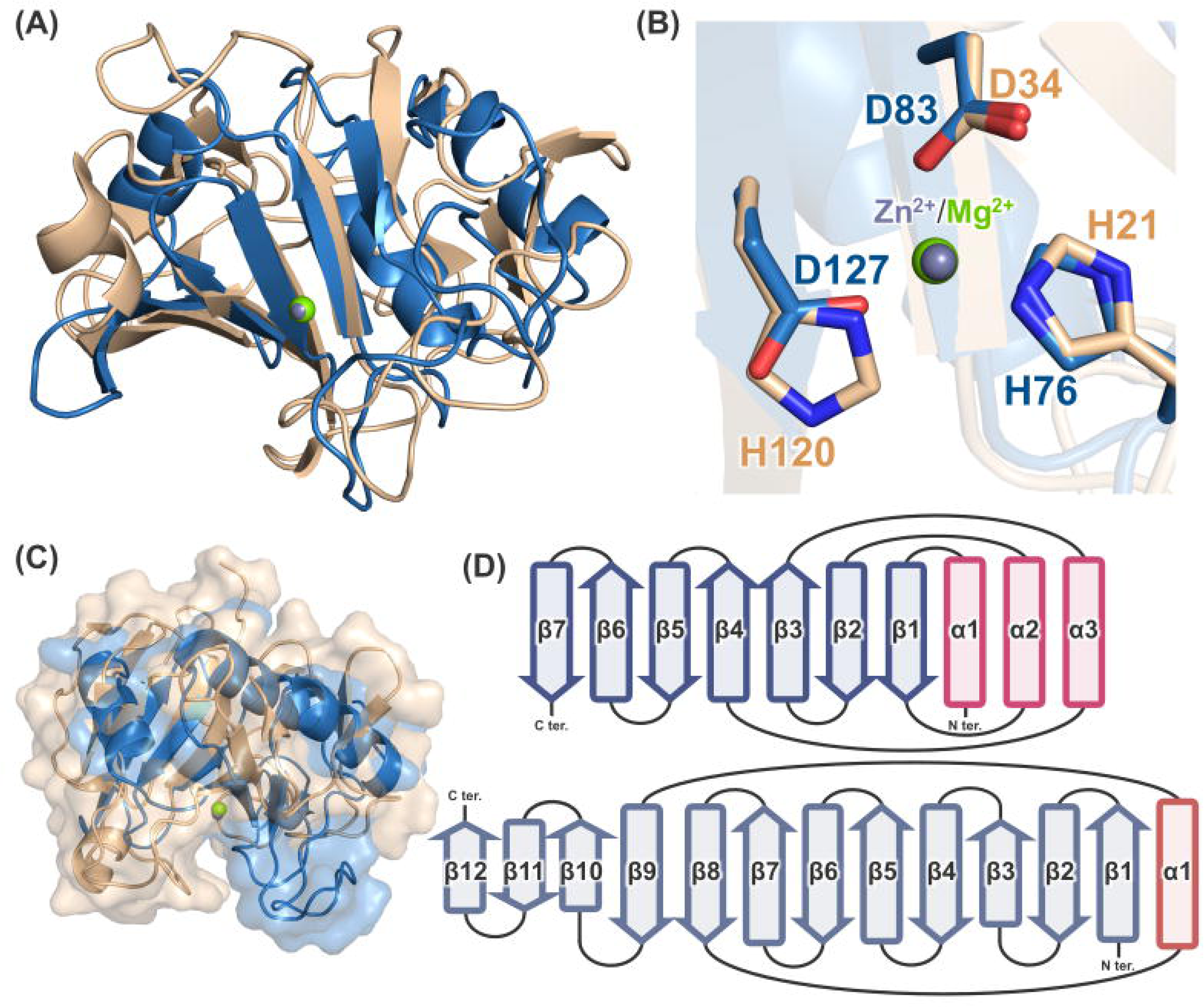
Structural comparison between *Rv*Peti and pseudoalterin. (A) Superposition of the structures of *Rv*Peti (blue) and pseudoalterin (beige), aligned based on their active-site regions. (B) Close-up view of the superimposed active sites. The metal ions bound in *Rv*Peti (Mg^2+^, light green) and in pseudoalterin (Zn^2+^, purple) are shown as spheres. (C) Surface representation of the superimposed structures, showing differences in the shape of the substrate-binding pocket between *Rv*Peti and pseudoalterin. (D) Topology diagrams of the secondary structure elements of *Rv*Peti (top) and pseudoalterin (bottom).

To further investigate the function of *Rv*Peti, we employed CLEAN, a tool that predicts enzyme function from amino acid sequence alone (Yu et al., 2023). CLEAN assigned *Rv*Peti to EC 3.5.1.28, classifying it as an N-acetylmuramyl-L-alanine amidase. A well-characterized example of this enzyme class is Rv3717 from *Mycobacterium tuberculosis*, which exhibits zinc-dependent amidase activity toward muramyl dipeptide (MDP), a model substrate of peptidoglycan (Prigozhin et al., 2013).

### 2.3. Binding of divalent metals alters the active-site environment

Given that both pseudoalterin and Rv3717 require zinc for catalysis, we investigated whether Zn^2+^ or Mn^2+^ could bind at the metal-binding site of *Rv*Peti. The *Rv*Peti crystals obtained were soaked in solutions containing ZnCl_2_ or MnCl_2_, and anomalous diffraction data were collected from the crystals. The resulting anomalous maps showed that Zn^2+^ and Mn^2+^ can occupy the metal-binding site formed by the HDD motif in *Rv*Peti (Figs. 3A, 3B, S1).

**Fig. 3.**
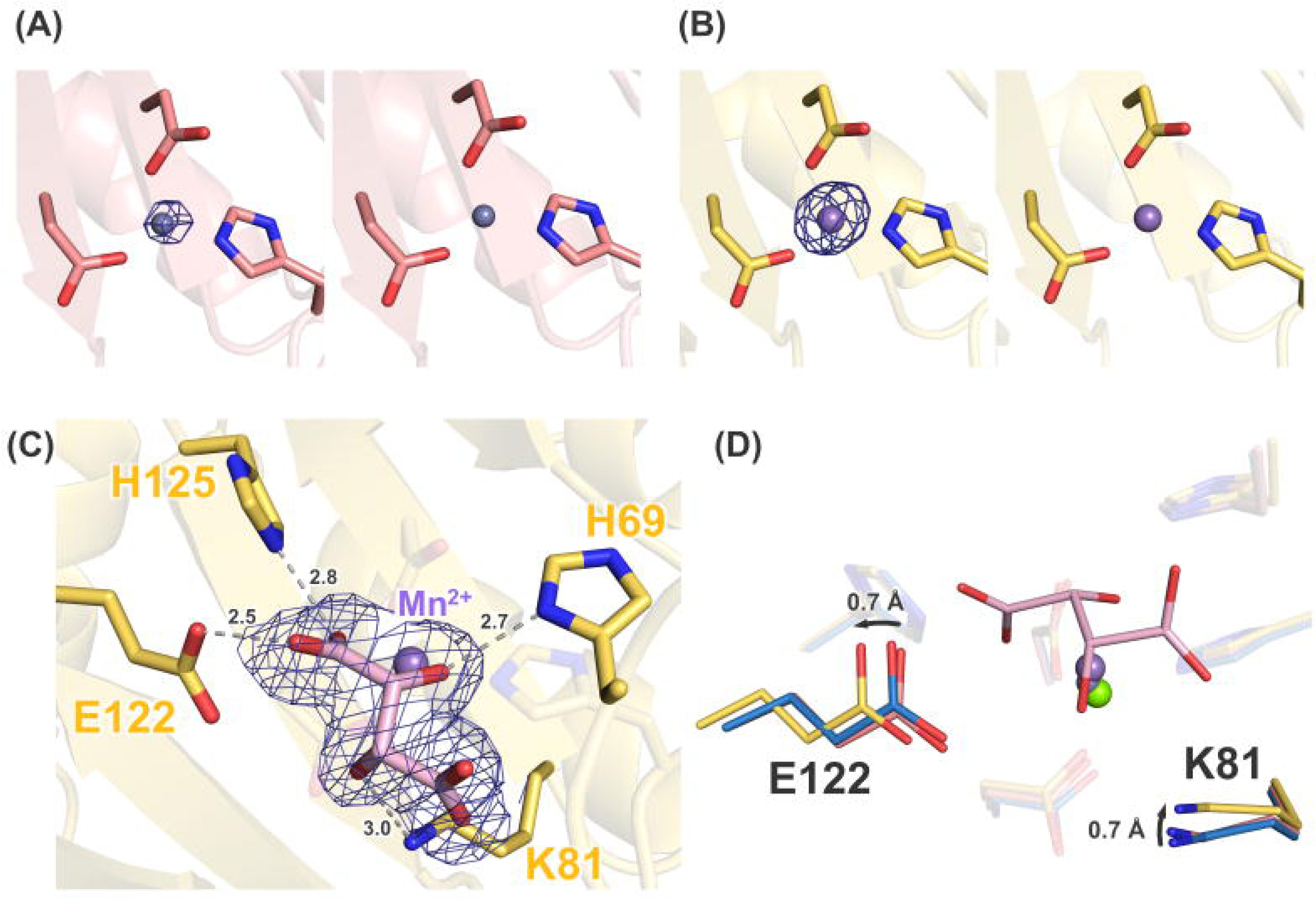
Identification of metal-binding sites in *Rv*Peti by anomalous scattering. (A) Anomalous scattering signal from the *Rv*Peti-Zn dataset. The anomalous peak is shown as a navy mesh contoured at 8.0σ. Diffraction data were collected at 1.28 Å (left) and 1.30 Å (right), respectively. (B) Anomalous scattering signal from the *Rv*Peti-Mn dataset. The anomalous peak is shown as a navy mesh contoured at 10.0σ. Diffraction data were collected at 1.80 Å (left) and 1.90 Å (right), respectively. (C) Close-up view of the active site in the Mn-soaked crystal structure, showing a tartrate molecule (pink, stick representation) near the active site, together with its corresponding electron density (2*F*_o_-*F*_c_ map contoured at 1.0σ). (D) Superposition of the apo (purified), Zn-soaked, and Mn-soaked structures of *Rv*Peti. In the *Rv*Peti-Mn, Lys81 and Glu122 show slight positional shifts relative to their positions in the other structures. A tartrate ion was observed near the active site of only the *Rv*Peti-Mn structure.

The overall structures did not show substantial differences compared to the as-purified form. However, in a crystal soaked with MnCl_2_, coordination of tartaric acid, which is a component of the crystallization solution, was observed (Fig. 3C). This tartaric acid interacts with His69, Lys81, Glu122, and His125 in the vicinity of the HDD motif. Consequently, Lys81 and Glu122 show slight conformational changes of approximately 0.7 Å compared to the as-purified structure (Fig. 3D), suggesting that these residues possess flexibility and may contribute to the stability of the bound ligand. Tartrate coordination was not observed in the absence of Mn^2+^ soaking, suggesting that Mn^2+^ binding alters the local ligand-binding environment. This observation raises the possibility that ligand binding depends on the identity of the bound metal ion.

Furthermore, in the Zn-bound crystal, several distances between the metal ion, the two water molecules, and surrounding amino acid residues in the hydrogen-bond network are shorter than in the as-purified structure (Fig. 4, S2). This suggests that the coordinated metal alters the local environment around the active site, which may influence catalysis.

**Fig. 4.**
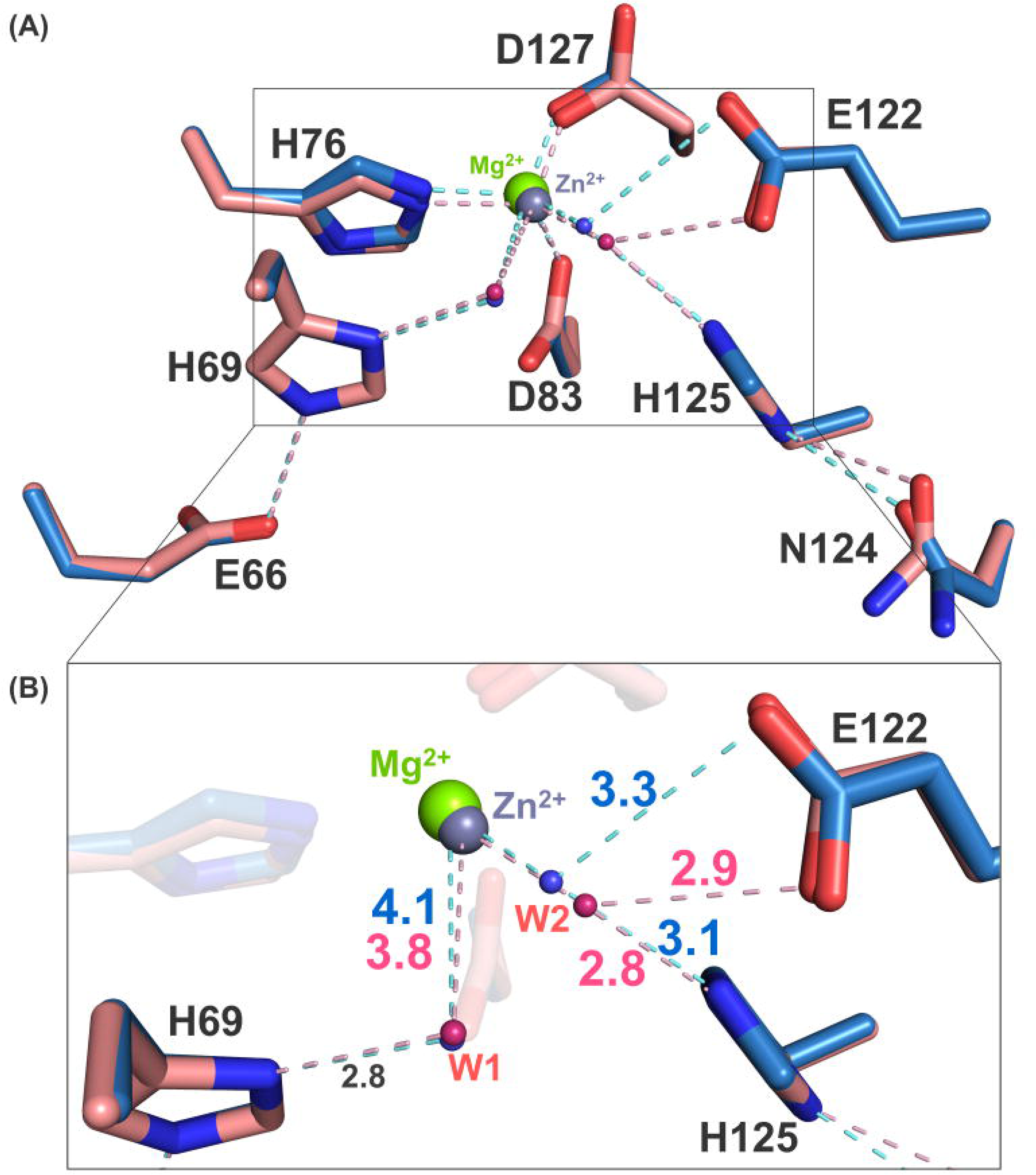
Metal-dependent changes in the hydrogen-bond network. (A) Superposition of the purified *Rv*Peti and *Rv*Peti-Zn structures. The as-purified structure is shown in blue, and the *Rv*Peti-Zn structure in pink. (B) Close-up view of the region showing the most pronounced change in hydrogen-bond distances upon Zn^2+^ binding, involving the metal-coordinated water molecules W1 (associated with His69) and W2 (associated with Glu122/His125).

### 2.4. Zn-dependent protease activity of *Rv*Peti

First, based on our structural study showing structural similarity of *Rv*Peti to pseudoalterin, we examined whether *Rv*Peti shows protease-mediated degradation of elastin. However, no reaction was observed even after 3 hours of incubation (Fig. S3). Similarly, *Rv*Peti showed no reactivity toward collagen, a polymeric peptide, which is also degraded by pseudoalterin. Additionally, we performed an amidase activity assay of *Rv*Peti using MDP, which is a possible substrate predicted by CLEAN, but no significant reaction was detected (Fig. S4).

Therefore, to detect general proteolytic activity, we performed a fluorescein isothiocyanate-labeled casein (FITC-casein) cleavage assay. In this assay, FITC-casein exists in a self-quenched state due to the proximity of the FITC groups. Proteolytic cleavage of the substrate increases the distance between the FITC groups, resulting in a rise in fluorescence intensity.

In the absence of added metal ions, the addition of *Rv*Peti resulted in a further decrease in fluorescence intensity (Fig. 5A). This may reflect binding of FITC-casein to the large groove of *Rv*Peti, which could shorten the distance between FITC groups and enhance self-quenching. No proteolytic activity toward FITC-casein was detected under this condition.

**Fig. 5.**
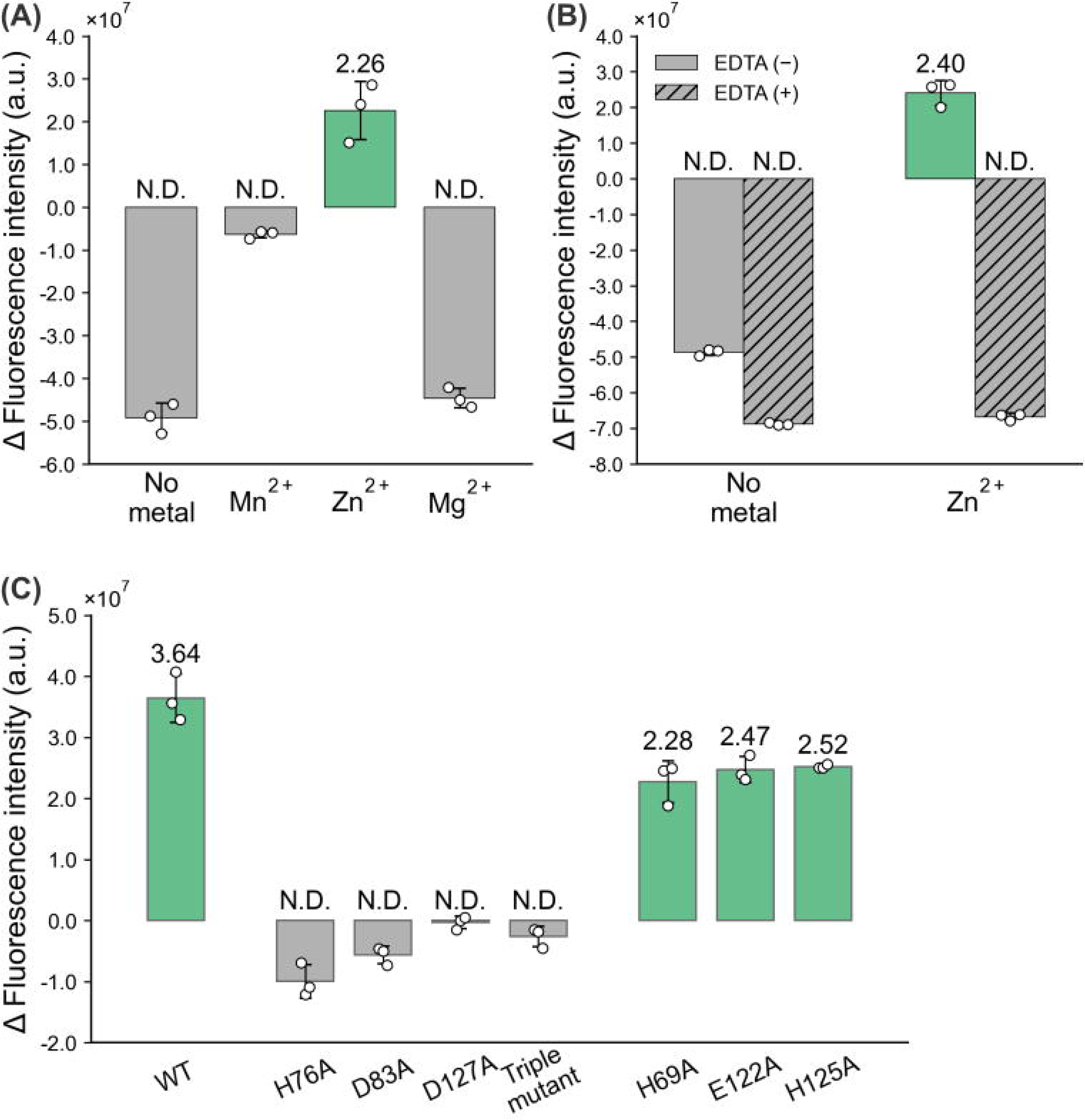
Zn^2+^-dependent proteolytic activity of *Rv*Peti. (A) Proteolytic activity of wild-type *Rv*Peti toward FITC-casein, measured in the absence of metal ions (No metal) or in the presence of Mn^2+^, Zn^2+^, or Mg^2+^. Activities of data with a negative Δ Fluorescence intensity are denoted as N.D. (not detected). Data is the mean ± SD; n = 3. (B) Effect of EDTA on proteolytic activity under no metal and Zn^2+^ conditions. Samples containing EDTA are indicated by hatched bars. (C) Proteolytic activity of wild-type and the indicated mutants (H76A, D83A, D127A, H76A/D83A/D127A triple mutant, H69A, E122A, and H125A) under Zn^2+^-supplemented conditions. Data is the mean ± SD; n = 3.

In contrast, Zn^2+^ supplementation led to a significant increase in fluorescence intensity relative to controls without added metal ions. This result indicates that Zn^2+^ supports the proteolytic activity of *Rv*Peti toward FITC-casein. On the other hand, protease activity was not observed in the presence of Mn^2+^ and Mg^2+^. Furthermore, chelation of Zn^2+^ by ethylenediaminetetraacetic acid (EDTA) significantly reduced this activity (Fig. 5B), strongly suggesting that *Rv*Peti functions as a Zn^2+^-dependent metalloprotease.

To further investigate the catalytic mechanism, we generated alanine substitution mutants of the HDD motif (H76A, D83A, D127A, and the triple mutant H76A/D83A/D127A) and residues surrounding the active site (H69A, E122A, and H125A) and evaluated their proteolytic activity using FITC-casein as a substrate in the presence of Zn^2+^ (Fig. 5C). Substitution of the zinc-coordinating residues His76, Asp83, and Asp127 markedly reduced proteolytic activity, supporting the importance of the zinc-coordinating site for proteolytic activity. In contrast, H69A, E122A, and H125A mutants retained approximately two-thirds of the wild-type activity. Because these residues form hydrogen bonds with water molecules at the active site, their substitution could alter the positions and/or stability of these water molecules and thereby affect catalysis. However, the effects of alanine substitution of these residues were modest. Therefore, whether these residues are primarily required for substrate stabilization or for the catalytic reaction itself remains an open question. Also, a detailed catalytic mechanism cannot be inferred from the current study.

Because pseudoalterin has been reported to cleave Gly-Gly bonds, we measured the peptidase activity using Gly-Gly-Gly (3Gly) as a substrate to examine whether *Rv*Peti can cleave this peptide substrate. In this reaction, the free amino groups generated by cleavage were detected using *o*-phthalaldehyde (OPA) in the presence of a reducing agent. No Zn-dependent difference in fluorescence was observed, indicating that efficient cleavage of 3Gly was not detected under the conditions tested (Fig. S6).

### 2.5. Weak esterase activity of *Rv*Peti

Since *Rv*Peti was suggested to possess protease activity, we investigated whether it also exhibits hydrolytic activity toward ester bonds, which similarly contain a carbonyl group. Esterase activity was assessed using *p*-nitrophenyl acetate (*p*NPA) as a substrate by measuring the absorbance of the released *p*-nitrophenol at 405 nm (Fig. 6A). Comparable increases in activity were observed in the presence of Mn^2+^ and Mg^2+^, as well as without added metal ions. In contrast, the reaction with Zn^2+^ yielded the lowest activity, which differed from the results obtained in the protease assay.

**Fig. 6.**
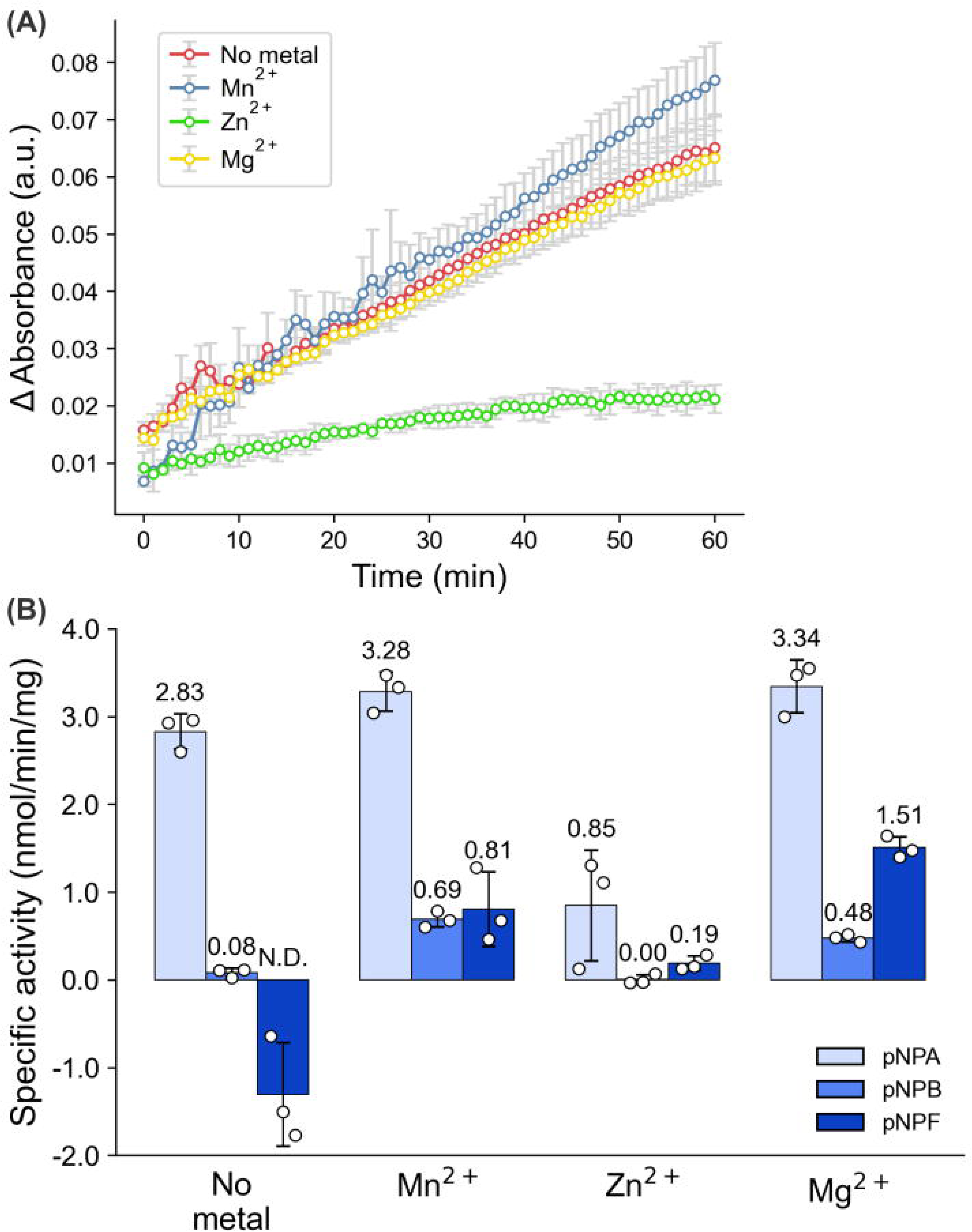
Esterase activity of *Rv*Peti. (A) Time course of esterase activity toward *p*NPA in the absence of metal ions (no metal, red) or in the presence of Mn^2+^ (blue), Zn^2+^ (green), or Mg^2+^ (yellow). Absorbance at 405 nm was measured every 1 min over a 60 min period. (B) Specific esterase activity of *Rv*Peti toward *p*NPA (light blue), *p*NPB (medium blue), and *p*NPF (dark blue), measured over 60min under No metal, Mn^2+^, Zn^2+^, or Mg^2+^ conditions. Activity is expressed as nmol/min/mg of enzyme.

To further investigate the substrate-recognition properties, we evaluated the hydrolytic activity toward *p*-nitrophenyl butyrate (*p*NPB), which has a longer acyl chain, and *p*-nitrophenyl trans-ferulate (*p*NPF), which has a bulkier aromatic group. Neither *p*NPB nor *p*NPF showed detectable activity under Zn^2+^ conditions, which had supported protease activity, whereas activity was observed under Mn^2+^ and Mg^2+^ conditions. The activity toward *p*NPB was less than one-fifth of that toward *p*NPA in all conditions. For *p*NPF, activity under Mg^2+^ conditions reached approximately half of that observed with *p*NPA, whereas under the other conditions, it remained as low as that observed with *p*NPB. These experiments show that the hydrolytic activity of *Rv*Peti varies among the tested substrates with different acyl chain lengths or side-chain structures.

Our results indicate that although *Rv*Peti exhibits hydrolytic activity toward *p*NPA, the activity is quite low. However, activity toward these model substrates alone does not establish whether *Rv*Peti functions physiologically as an esterase. In fact, the specific esterase activity of *Rv*Peti was only on the order of a few nmol/min/mg of enzyme (Fig. 6B). The purified porcine liver carboxylesterases PLE1 and PL6 exhibit specific activities of 1,700 and 2,130 nmol/min/mg (Zhou et al., 2019), respectively, whereas recombinant *Pseudomonas fluorescens* esterase (PEE) exhibits 120,000 nmol/min/mg (Krebsfänger et al., 1998). Thus, the activity of *Rv*Peti is substantially lower than those reported for representative esterases. Hydrolysis of ester bonds has previously been reported even with a minimal catalytic system consisting of Glu-His-water or His-His-water (Novak et al., 2013; Khersonsky & Tawfik, 2006). Therefore, it is possible that, upon binding of *p*NPA within the active pocket of *Rv*Peti, a water molecule is appropriately positioned to facilitate incidental hydrolysis of the ester bonds. Thus, the observed esterase activity may represent a secondary activity arising from substrate binding within the pocket, although its physiological relevance remains to be established.

## 3. Discussion

In this study, we determined the crystal structure of *Rv*Peti from the tardigrade *R. varieornatus* and characterized its basic functional properties. Proteins with similar amino acid sequences are found in many bacteria and other eukaryotes, suggesting that this gene is broadly distributed. Because this protein family is conserved across a wide range of organisms, it is likely to have physiological significance; however, its molecular properties, including its enzymatic activity, have remained unknown. We found that *Rv*Peti adopts an overall structure distinct from those of previously characterized proteins and exhibits Zn^2+^-dependent protease activity. These findings suggest that *Rv*Peti belongs to a novel protease family and provide an important foundation for elucidating its structural and functional properties.

*Rv*Peti differs from many other proteases, including the M23 proteases such as pseudoalterin, by its size and simple domain architecture. With a molecular weight of approximately 14 kDa, *Rv*Peti is close in size to the isolated catalytic domain of LytM, an M23 peptidase from *Staphylococcus aureus*, which has been shown to be sufficient to form the M23 peptidase fold and catalytic site (Firczuk et al., 2005). In M23 peptidases, the metal-binding and catalytic sites are generally located within a single domain, but the family typically also contains additional domains beyond this catalytic core (Razew et al., 2022). *Rv*Peti, by contrast, appears to comprise a compact single domain that contains both the metal-binding and putative catalytic sites.

Notably, the metal-binding residues are composed of a His-Asp-Asp combination, a feature shared among homologous proteins identified through BLAST searches in other organisms. In contrast, the HHD motif is well known as the catalytic center of a broad range of metal-dependent hydrolases, including metalloproteases, amidases, and deacetylases. The significance of *Rv*Peti employing an HDD motif rather than the canonical HHD motif remains unclear. It is currently not possible to determine whether this substitution merely reflects a phylogenetic bias or instead points to a catalytic mechanism distinct from that of other metalloproteases. Further functional analyses will be needed to determine the extent of the proteolytic activity that *Rv*Peti exhibits toward its native substrate.

*Rv*Peti was confirmed to exhibit Zn^2+^-dependent proteolytic activity toward casein. However, this activity was more than an order of magnitude lower than that of trypsin, which was assayed in parallel as a positive control (Fig. S5). One possibility is that the relatively open substrate-binding groove of *Rv*Peti affects substrate positioning, such that a substrate of appropriate size and shape may be required for efficient cleavage. Similarly, pseudoalterin shows high reactivity toward elastin but low reactivity toward collagen and casein. The difference in reactivity between *Rv*Peti and trypsin may reflect differences in substrate preference, and casein may not be an optimal substrate for *Rv*Peti. Although the true physiological substrate of *Rv*Peti could not be identified in this study, elucidating its precise substrate range will be an important focus of future research.

We also identified two water molecules near the active center whose positions changed slightly in the Zn^2+^-bound crystal structure. The water molecule at the catalytic site of metalloproteases is activated by a metal ion and often performs a nucleophilic attack on the carbonyl carbon atom of the substrate. Our current study could not determine whether the two water molecules observed at the catalytic site in *Rv*Peti serve as important catalytic components. Alanine mutants of His69, Glu122, and His125, residues that interact with these water molecules, each retained approximately two-thirds of the wild-type activity, but these results do not resolve the individual catalytic roles of the two water molecules. Whether either water molecule participates in nucleophilic attack may depend on the orientation in which the substrate binds within the active-site pocket. Resolving the detailed catalytic mechanism of *Rv*Peti would benefit from identifying a suitable substrate and subsequently obtaining a substrate-complex structure.

Weak but detectable esterase activity was also observed. This activity varied depending on the metal condition. Zn^2+^, which supported protease activity in our assays, was associated with lower esterase activity. This suggests that the identity of the metal ion bound at the active site may alter the catalytic environment and/or substrate binding. Accordingly, the local availability of metal ions and/or metallochaperone proteins could influence the reactivity of *Rv*Peti *in vivo*, although this possibility remains to be tested. How *Rv*Peti is actually regulated and utilized within the tardigrade, and what physiological role it plays *in vivo*, remain unknown at present.

This study reveals the existence of a novel protease family that, despite being present across a wide range of organisms, has been largely overlooked until now, representing a first step toward fully characterizing this family. In particular, *Rv*Peti possesses a compact single-domain architecture that accommodates both metal binding and proteolytic activity. From an evolutionary perspective, the gene encoding *Rv*Peti has been proposed to have been acquired through HGT. The broader question of how horizontally acquired genes may contribute to the acquisition of host stress tolerance remains open to debate. Interestingly, several organisms identified through the sequence homology search of *Rv*Peti possess some form of environmental tolerance capability. For example, *T. hazakensis* is a thermophilic bacterium (Yabe et al., 2010), and *Folsomia candida* is a springtail that shows physiological adaptation to dry soil environments through osmotic regulation (Sjursen et al., 2001). Although these observations do not establish a direct relationship between the Peti family and environmental stress tolerance, they motivate further investigation of the physiological roles of this protein family in different organisms. Further characterization of Peti family proteins and identification of their native substrates will be important for clarifying their evolutionary and physiological significance.

## 4. Methods

### 4.1. Databases and sequence analyses

Protein sequences of interest were obtained from the database of the National Center for Biotechnology Information (https://www.ncbi.nlm.nih.gov). Amino acid sequence analyses were performed using MEGA11 (Tamura et al., 2021). BLAST searches were performed via the web service for protein BLAST (https://blast.ncbi.nlm.nih.gov/Blast.cgi) (Altschul et al., 1990). A maximum likelihood method was used to estimate phylogenetic trees. Phylogenetic trees were visualized using iTOL (Letunic & Bork, 2021). Multiple sequence alignment was performed using Clustal Omega (https://www.ebi.ac.uk/jdispatcher/msa/clustalo) (Sievers & Higgins, 2014) and visualized by ESPript 3 (http://espript.ibcp.fr) (Robert & Gouet, 2014). Pairwise sequence similarity between *Rv*Peti and pseudoalterin was calculated using EMBOSS Needle with the BLOSUM62 substitution matrix (https://www.ebi.ac.uk/jdispatcher/psa/emboss_needle) (Rice et al., 2000). Model structures predicted by AlphaFold2 (as of Apr 2025) were obtained from AlphaFold Protein Structure Database (https://alphafold.ebi.ac.uk/).

### 4.2. Cloning of *Rv*Peti

An expression plasmid for *Rv*Peti expression in *E. coli* was constructed. The RvY_14254 gene was obtained as a synthetic DNA fragment from Integrated DNA Technologies (IDT, Coralville, IA, USA). The pET-28a (+) vector (Novagen, Madison, WI, USA) was linearized and amplified by PCR using the primers listed in Table. S2. The DNA fragment containing the RvY_14254 gene was cloned into the linearized pET-28a (+) vector using In-Fusion Snap Assembly Master Mix (Takara Bio). All DNA sequences were checked by DNA sequencing (Fasmac Co., Ltd., Atsugi, Kanagawa, Japan).

### 4.3. Expression and purification

The expression construct of *Rv*Peti contains an N-terminal 6×His tag followed by a TEV protease recognition site. The final amino acid sequence is as follows, in which italic characters indicate the *Rv*Peti region and ENLYFQS is a TEV protease recognition site.

*MGSSHHHHHHENLYFQSRLTHAQALQRVNAVGISVHSSGNCSDRNRPNCTSLDTILAKTIDQIVAFKRKSGKVVIITGGTETGHASGTKSHWNGYKLDIAMNAGVTDYITKNFRYIGTTNGGRWAQYQDGSDIWTNEGNHW DIVIS*

Site-directed mutagenesis was performed by inverse PCR using the pET-28a (+)-*Rv*Peti plasmid as a template and the KOD -Plus-Mutagenesis Kit (TOYOBO, Osaka, Japan), generating the H69A, D83A, E122A, H125A, and D127A mutants. The primers used for each mutant are shown in Table S2. The triple mutant was generated using a synthetic DNA fragment obtained from IDT carrying alanine substitutions at positions 76, 83, and 127, which were inserted into the vector. All mutations were verified by DNA sequencing.

The pET-28a (+)-*Rv*Peti expression construct was transformed into *E. coli* Shuffle T7 (New England BioLabs, Ipswich, MA, USA). Transformed *E. coli* cells were grown at 30 °C in LB medium (Merck, Darmstadt, Germany) with 50 µg/mL kanamycin sulfate (FUJIFILM Wako Pure Chemical Corporation, Osaka, Japan) until OD_600_ reached 1.0-1.2. The protein expression was induced by the addition of isopropyl β-D-1-thiogalactopyranoside (IPTG) (Nacalai Tesque, Nakagyo-ku, Kyoto, Japan) at a final concentration of 0.5 mM. After 20 h at 18 °C, cells were collected by centrifugation (6000 rpm, 30 min) at 4 °C and resuspended in Buffer A (20 mM Tris-HCl pH 8.0, 200 mM NaCl, and 5 mM imidazole). Cells were lysed by sonication on ice, and cell debris was removed by centrifugation (20000 *g*, 90 min) at 4 °C. The supernatant was loaded onto a HiTrap TALON column (GE Healthcare, Chicago, IL, USA) previously equilibrated with buffer A. The column was then washed with Buffer A and His-*Rv*Peti was eluted with Buffer B (20 mM Tris-HCl pH 8.0, 200 mM NaCl, and 200 mM imidazole). Preliminary attempts to remove the N-terminal His tag using TEV protease were unsuccessful. Therefore, we used His-*Rv*Peti throughout this study; that is, all protein preparations used for structural and biochemical analyses were purified without TEV protease treatment. Fractions containing His-*Rv*Peti, as confirmed by SDS-PAGE, were concentrated using a Amicon® Ultra Centrifugal Filter, 10 kDa MWCO (Merck Millipore, Darmstadt, Germany) and loaded onto a HiLoad™ 16/60 Superdex™ 75 pg size-exclusion chromatography (SEC) column (GE Healthcare) equilibrated with SEC buffer (20 mM Tris-HCl, pH 8.0, 150 mM NaCl). The eluted fractions were analyzed by SDS-PAGE, and fractions containing His-*Rv*Peti were collected (Fig. S7). The *Rv*Peti fraction was concentrated to approximately 25 mg/mL and stored at -80 °C until use.

### 4.4. Crystallization

Crystallization was performed by the sitting drop vapor-diffusion method. Crystals of *Rv*Peti appeared under the condition of 25 mg/ml protein, 0.2 M sodium tartrate dibasic dihydrate, 26-32 % w/v polyethylene glycol 3,350 at 20 °C. A crystal was transferred to the corresponding cryoprotectant solutions containing 5% (v/v) ethylene glycol and then flash-cooled in liquid nitrogen. For metal soaking, *Rv*Peti crystals were soaked for 1 min in the crystallization solution supplemented with 1 mM ZnCl or MnCl . The Zn-soaked and Mn-soaked crystals (*Rv*Peti–Zn and *Rv*Peti–Mn, respectively) were subsequently transferred to the corresponding cryoprotectant solutions containing 5% (v/v) ethylene glycol and 1 mM ZnCl or MnCl , respectively, and then flash-cooled in liquid nitrogen.

### 4.5. Data collection and structure determination

Crystals of *Rv*Peti were harvested using LithoLoops (WakenBtech Co.,Ltd, Sakyo-ku, Kyoto, Japan) and cryo-cooled in liquid nitrogen. X-ray diffraction data were collected at the BL44XU beamline of SPring-8 (Hyogo, Japan) using an EIGER X 16M detector (Dectris, Baden-Dättwil, Switzerland) at 100 K. The native dataset was collected at a wavelength of 0.9 Å. Anomalous diffraction data were collected to identify metal ions: at 1.28 Å and 1.30 Å for Zn and at 1.80 Å and 1.90 Å for Mn. Diffraction data were processed using XDS (Kabsch, 2010) and scaled with AIMLESS (Evans & Murshudov, 2013). The resolution cutoff was determined based on CC_1/2_ values. Phases were determined by the molecular replacement method using MOLREP in the CCP4 suite (Murshudov et al., 2011) with AlphaFold model as a search model. The model was manually adjusted using COOT (Emsley et al., 2010). REFMAC5 in the CCP4 suite and phenix.refine (Adams et al., 2010) in the PHENIX software suite were used for refinement. Metal ions were assigned based on coordination geometry, and the Zn and Mn assignment were confirmed by anomalous difference Fourier maps. The stereochemical quality of the final model was evaluated using MolProbity (Chen et al., 2010). All structure figures were prepared using PyMOL (The PyMOL Molecular Graphics System, Version 3.0 Schrödinger, LLC.).

Data collection and refinement statistics are summarized in Table 1. The coordinate and structure factor files are deposited at the Protein Data Bank (extended PDB code IDs: pdb_000045MZ; pdb_000045NE; pdb_000045NH).

**Table 1.** Data collection and refinement statistics.

|  | **RvPeti** | **RvPeti_Zn** | **RvPeti_Mn** |
| --- | --- | --- | --- |
| **Data collection** |  |  |  |
| **Diffraction source** | SPring-8 BL44XU |  |  |
| **Wavelength (Å)** | 0.9 |  |  |
| **Resolution range** | 49.46 - 1.35 (1.37 - 1.35) | 36.05 - 1.85 (1.89 - 1.85) | 41.28 - 1.80 (1.84 - 1.80) |
| **Space group** | *P* 3<sub>2</sub> 2 1 | *P* 3<sub>2</sub> 2 1 | *P* 3<sub>2</sub> 2 1 |
| ***a*, *b*, *c* (Å)** | 72.56, 72.56, 80.18 | 72.10, 72.10, 81.07 | 71.72, 71.72, 82.57 |
| ***α*, *β*, *γ* (°)** | 90, 90, 120 | 90, 90, 120 | 90, 90, 120 |
| **Total reflections** | 531906 | 140756 | 227851 |
| **Unique reflections** | 54058 (2635) | 21035 (1287) | 23263 (1367) |
| **Redundancy** | 9.8 (10.3) | 6.7 (6.9) | 9.8 (10.3) |
| **Completeness (%)** | 100.00 (100.00) | 99.3 (99.8) | 100.0 (100.0) |
| ***I*/σ (*I*)** | 17.6 (1.6) | 12.5 (1.3) | 14.8 (1.4) |
| ***R*<sub>merge</sub> (all *I*+ & *I*-)** | 0.052 (1.467) | 0.086 (1.652) | 0.076 (1.795) |
| ***R*<sub>meas</sub> (all *I*+ & *I*-)** | 0.055 (1.544) | 0.094 (1.783) | 0.080 (1.889) |
| ***R*<sub>pim</sub> (all *I*+ & *I*-)** | 0.018 (0.479) | 0.035 (0.659) | 0.026 (0.586) |
| **CC<sub>1/2</sub>** | 0.999 (0.729) | 0.998 (0.529) | 0.999 (0.574) |
| **Refinement** |  |  |  |
| **Resolution range** | 49.46 - 1.35 (1.37 - 1.35) | 36.05 - 1.85 (1.95 - 1.85) | 35.86 - 1.80 (1.88 - 1.80) |
| **No. of reflections, working set** | 54006 (2651) | 21030 (2976) | 23225 (2860) |
| **No. of reflections, test set** | 2754 (131) | 1035 (133) | 1128 (130) |
| **$R_{\text{work}}/R_{\text{free}}$ (%)** | 15.02/16.41<br>(28.72/29.40) | 17.17/18.79<br>(27.70/31.37) | 17.55/20.76<br>(28.38/28.28) |
| **No. of model atoms** |  |  |  |
| **macromolecules** | 1127 | 1041 | 1040 |
| **ligands** | 25 | 23 | 25 |
| **solvent** | 203 | 152 | 139 |
| **Protein residues** | 138 | 133 | 133 |
| **RMS bonds (Å)** | 0.004 | 0.006 | 0.005 |
| **RMS angles (°)** | 0.67 | 0.70 | 0.75 |
| **Ramachandran plot (%)** |  |  |  |
| **Favored** | 99.26 | 99.24 | 99.24 |
| **Allowed** | 0.74 | 0.76 | 0.76 |
| **Outliers** | 0.00 | 0.00 | 0.00 |
| **Average $B$-factor (Å<sup>2</sup>)** |  |  |  |
| **Overall** | 33.23 | 44.13 | 41.43 |
| **macromolecules** | 30.72 | 42.86 | 40.47 |
| **ligands** | 61.97 | 75.58 | 50.54 |
| **solvent** | 43.63 | 48.14 | 46.99 |
| **PDB code ID** | pdb_000045MZ | pdb_000045NE | pdb_000045NH |

Statistics for the highest-resolution shell are shown in parentheses.

### 4.6. Enzymatic assay

Proteolytic activity toward elastin and collagen was assessed using elastin and collagen, respectively, as substrates (Zhao et al., 2012). Reactions were performed with 5 mg/mL substrate and 4 μg *Rv*Peti in 20 mM Tris-HCl (pH 8.0), 150 mM NaCl, and 10 μM metal ion. Reaction solutions were incubated at 37 °C with continuous stirring for 3 h. Substrates incubated without *Rv*Peti served as negative controls. Substrate degradation was assessed by visual inspection and documented by photographing the samples at regular intervals during incubation.

Protease activity was measured using FITC-casein from bovine milk (Type III, essentially salt-free, lyophilized powder; Sigma-Aldrich) as the substrate. Reactions were performed in 20 mM Tris-HCl (pH 8.0), 150 mM NaCl, containing 100 μg/mL FITC-casein and 5.0 μM purified *Rv*Peti, in the presence or absence of 5 μM MnCl , ZnCl , or MgCl . To examine the metal dependence of the proteolytic activity, reactions were also performed in the presence of 1 mM EDTA. Reactions were incubated at 37 °C for 60 min. Fluorescence was measured at excitation/emission wavelengths of 490/525 nm using a SpectraMax iD3s multi-mode microplate reader (Molecular Devices, San Jose, CA, USA). The fluorescence difference was calculated by subtracting the fluorescence intensity of the corresponding enzyme-free control from that of the enzyme-containing reaction after 60 min. Measurements were performed in triplicate using aliquots prepared from a single reaction mixture, and data are presented as mean ± SD.

Amidase activity toward MDP and peptidase activity toward 3Gly were measured using an OPA assay, which detects free amino groups generated upon substrate cleavage (Prigozhin et al., 2013). Reactions were performed in 50 mM sodium phosphate buffer containing 500 μM MDP or 3Gly and 5 μM *Rv*Peti, with or without 10 μM metal ion. Reactions were incubated at 37 °C for 60 min, and *Rv*Peti was removed by centrifugation through a 3-kDa molecular weight cutoff filter prior to the OPA assay to minimize the contribution of protein N-termini to the OPA signal. A 20-μL aliquot of each sample was mixed with 100 μL of OPA developing solution (1.5 mM *o*-phthalaldehyde, 350 mM sodium borate, pH 9.5, and 1% (v/v) β-mercaptoethanol), and fluorescence was measured at excitation/emission wavelengths of 338/455 nm.

Esterase activity was measured using *p*NPA, *p*NPB, and *p*NPF as substrates. Reactions were performed in 20 mM Tris-HCl (pH 7.5), 150 mM NaCl, containing 1 mM substrate and 10 μM *Rv*Peti, with or without 100 μM metal ion. *p*NPF was first dissolved in DMSO, followed by the addition of Triton X-100 to a final concentration of 2.5%. Reactions were incubated at 37 °C for 60 min, and substrate hydrolysis was monitored by measuring the absorbance of the released *p*NP at 405 nm. The change in absorbance was calculated from the difference between the measurements at 60 and 0 min, after subtraction of the corresponding enzyme-free control.

## Supplement information

Table S1. Data collection of anomalous data.

Table S2. Primers used in this research (from 5’ to 3’).

Fig. S1. Close-up view of *Rv*Peti-Zn and *Rv*Peti-Mn.

Fig. S2. Detailed hydrogen-bond network surrounding the Zn^2+^-binding site in *Rv*Peti-Zn.

Fig. S3. *Rv*Peti does not hydrolyze elastin or collagen.

Fig. S4. OPA assay for MDP cleavage by *Rv*Peti.

Fig. S5. Comparison of Zn^2+^-dependent proteolytic activity of *Rv*Peti with trypsin.

Fig. S6. Peptidase activity of *Rv*Peti toward Gly-Gly-Gly.

Fig. S7 Results of size exclusion chromatography and SDS-PAGE of *Rv*Peti.

## Supplement data

Data S1. Sequences used for alignment (supporting Fig. 1A)

## AUTHOR CONTRIBUTIONS

**Tomoka Higashi**: conceptualization, investigation, writing—original draft, writing—review and editing, formal analysis, data curation, visualization, validation. **Yohta Fukuda**: conceptualization, investigation, funding acquisition, writing—original draft, writing—review and editing, formal analysis, supervision, project administration, data curation, validation. **Tsuyoshi Inoue**: funding acquisition, supervision.

## ACKNOWLEDGMENTS

X-ray data were collected at SPring-8 BL44XU under the Collaborative Research Program of the Institute for Protein Research, Osaka University (Proposal Nos. 2025A6524, 2025B6524, 2026A6923). We acknowledge support from beamline staff.

The authors used ChatGPT and Claude to assist with English language editing and grammatical improvement. All AI-assisted text was reviewed and edited by the authors, who take full responsibility for the final content of the manuscript.

## DATA AVAILABILITY STATEMENT

The data that support the findings of this study are openly available in the Protein Data Bank at https://www.rcsb.org/, reference numbers pdb_000045MZ (as-purified structure); pdb_000045NE (Zn-bound state); pdb_000045NH (Mn-bound state). The raw X-ray diffraction data collected in this study are available at the Xtal Raw Data Archive (https://xrda.pdbj.org) under the IDs corresponding to the Protein Data Bank depositions.

## References

Adams PD, Afonine PV, Bunkóczi G, Chen VB, Davis IW, Echols N, et al. PHENIX: a comprehensive Python-based system for macromolecular structure solution. Acta Cryst D. 2010;66(2):213–21. 10.1107/S0907444909052925

Altschul SF, Gish W, Miller W, Myers EW, Lipman DJ. Basic local alignment search tool. Journal of Molecular Biology. 1990;215(3):403–10. 10.1016/S0022-2836(05)80360-2

Becquerel P. La Suspension De La Vie Au-Dessous De 1/20-Degrees-K Absolu Par Demagnetisation Adiabatique De Lalun De Fer Dans Le Vide Le Plus Eleve. C R Hebd Seances Acad Sci. 1950;231(4):261–3.

Boothby TC, Tapia H, Brozena AH, Piszkiewicz S, Smith AE, Giovannini I, et al. Tardigrades Use Intrinsically Disordered Proteins to Survive Desiccation. Molecular Cell. 2017;65(6):975–984.e5. 10.1016/j.molcel.2017.02.018

Boothby TC, Tenlen JR, Smith FW, Wang JR, Patanella KA, Osborne Nishimura E, et al. Evidence for extensive horizontal gene transfer from the draft genome of a tardigrade. Proceedings of the National Academy of Sciences. 2015;112(52):15976–81. 10.1073/pnas.1510461112

Chen VB, Arendall WB, Headd JJ, Keedy DA, Immormino RM, Kapral GJ, et al. MolProbity: all-atom structure validation for macromolecular crystallography. Acta Cryst D. 2010;66(1):12–21. 10.1107/S0907444909042073

Emsley P, Lohkamp B, Scott WG, Cowtan K. Features and development of Coot. Acta Cryst D. 2010;66(4):486–501. 10.1107/S0907444910007493

Evans PR, Murshudov GN. How good are my data and what is the resolution? Acta Cryst D. 2013;69(7):1204–14. 10.1107/S0907444913000061

Firczuk M, Mucha A, Bochtler M. Crystal Structures of Active LytM. Journal of Molecular Biology. 2005;354(3):578–90. 10.1016/j.jmb.2005.09.082

Fukuda Y, Miura Y, Mizohata E, Inoue T. Structural insights into a secretory abundant heat-soluble protein from an anhydrobiotic tardigrade, Ramazzottius varieornatus. FEBS Letters. 2017;591(16):2458–69. 10.1002/1873-3468.12752

Guidetti R, Altiero T, Rebecchi L. On dormancy strategies in tardigrades. Journal of Insect Physiology. 2011;57(5):567–76. 10.1016/j.jinsphys.2011.03.003

Hashimoto T, Horikawa DD, Saito Y, Kuwahara H, Kozuka-Hata H, Shin-I T, et al. Extremotolerant tardigrade genome and improved radiotolerance of human cultured cells by tardigrade-unique protein. Nat Commun. 2016;7(1):12808. 10.1038/ncomms12808

Hengherr S, Worland MR, Reuner A, Brümmer F, Schill RO. High-Temperature Tolerance in Anhydrobiotic Tardigrades Is Limited by Glass Transition. Physiological and Biochemical Zoology. 2009;82(6):749–55. 10.1086/605954

Holm L. Dali server: structural unification of protein families. Nucleic Acids Res. 2022;50(W1):W210–5. 10.1093/nar/gkac387

Horikawa DD, Cumbers J, Sakakibara I, Rogoff D, Leuko S, Harnoto R, et al. Analysis of DNA Repair and Protection in the Tardigrade Ramazzottius varieornatus and Hypsibius dujardini after Exposure to UVC Radiation. PLOS ONE. 2013;8(6):e64793. 10.1371/journal.pone.0064793

Horikawa DD, Sakashita T, Katagiri C, Watanabe M, Kikawada T, Nakahara Y, et al. Radiation tolerance in the tardigrade Milnesium tardigradum. International Journal of Radiation Biology. 2006;82(12):843–8. 10.1080/09553000600972956

Kabsch W. XDS. Acta Cryst D. 2010;66(2):125–32. 10.1107/S0907444909047337

Kamilari M, Pust FL, Marcantognini A, Møbjerg A, Neves RC, Schiøtt M, et al. At the Edge of Survival: Exploring the Frontiers of Tardigrade Extreme Stress Tolerance. Molecular Ecology. 2026;35(14):e70471. 10.1111/mec.70471

Kasianchuk N, Rzymski P, Kaczmarek Ł. The biomedical potential of tardigrade proteins: A review. Biomedicine & Pharmacotherapy. 2023;158:114063. 10.1016/j.biopha.2022.114063

Kato S, Deguchi K, Obana M, Fujio Y, Fukuda Y, Inoue T. Metabolite phosphatase from anhydrobiotic tardigrades. The FEBS Journal. 2024;291(23):5195–213. 10.1111/febs.17296

van Kempen M, Kim SS, Tumescheit C, Mirdita M, Lee J, Gilchrist CLM, et al. Fast and accurate protein structure search with Foldseek. Nat Biotechnol. 2024;42(2):243–6. 10.1038/s41587-023-01773-0

Khersonsky O, Tawfik DS. The Histidine 115-Histidine 134 Dyad Mediates the Lactonase Activity of Mammalian Serum Paraoxonases *. Journal of Biological Chemistry. 2006;281(11):7649–56. 10.1074/jbc.M512594200

Koutsovoulos G, Kumar S, Laetsch DR, Stevens L, Daub J, Conlon C, et al. No evidence for extensive horizontal gene transfer in the genome of the tardigrade Hypsibius dujardini. Proceedings of the National Academy of Sciences. 2016;113(18):5053–8. 10.1073/pnas.1600338113

Krebsfänger N, Zocher F, Altenbuchner J, Bornscheuer UT. Characterization and enantioselectivity of a recombinant esterase from Pseudomonas fluorescens. Enzyme and Microbial Technology. 1998;22(7):641–6. 10.1016/S0141-0229(98)00004-0

Letunic I, Bork P. Interactive Tree Of Life (iTOL) v5: an online tool for phylogenetic tree display and annotation. Nucleic Acids Res. 2021;49(W1):W293–6. 10.1093/nar/gkab301

Li Lei, Ge Z, Liu S, Zheng K, Li Y, Chen K, et al. Multi-omics landscape and molecular basis of radiation tolerance in a tardigrade. Science. 2024;386(6720):eadl0799. 10.1126/science.adl0799

Lim S, Reilly CB, Barghouti Z, Marelli B, Way JC, Silver PA. Tardigrade secretory proteins protect biological structures from desiccation. Commun Biol. 2024;7(1):633. 10.1038/s42003-024-06336-w

Møbjerg N, Neves RC. New insights into survival strategies of tardigrades. Comparative Biochemistry and Physiology Part A: Molecular & Integrative Physiology. 2021;254:110890. 10.1016/j.cbpa.2020.110890

Murshudov GN, Skubák P, Lebedev AA, Pannu NS, Steiner RA, Nicholls RA, et al. REFMAC5 for the refinement of macromolecular crystal structures. Acta Cryst D. 2011;67(4):355–67. 10.1107/S0907444911001314

Nelson DR. Current Status of the Tardigrada: Evolution and Ecology. Integr Comp Biol. 2002;42(3):652–9. 10.1093/icb/42.3.652

Novak HR, Sayer C, Isupov MN, Paszkiewicz K, Gotz D, Mearns Spragg A, et al. Marine Rhodobacteraceae l-haloacid dehalogenase contains a novel His/Glu dyad that could activate the catalytic water. The FEBS Journal. 2013;280(7):1664–80. 10.1111/febs.12177

Ono F, Saigusa M, Uozumi T, Matsushima Y, Ikeda H, Saini NL, et al. Effect of high hydrostatic pressure on to life of the tiny animal tardigrade. Journal of Physics and Chemistry of Solids. 2008;69(9):2297–300. 10.1016/j.jpcs.2008.04.019

Prigozhin DM, Mavrici D, Huizar JP, Vansell HJ, Alber T. Structural and Biochemical Analyses of Mycobacterium tuberculosis N-Acetylmuramyl-l-alanine Amidase Rv3717 Point to a Role in Peptidoglycan Fragment Recycling*. Journal of Biological Chemistry. 2013;288(44):31549–55. 10.1074/jbc.M113.510792

Razew A, Schwarz J-N, Mitkowski P, Sabala I, Kaus-Drobek M. One fold, many functions—M23 family of peptidoglycan hydrolases. Front Microbiol. 2022;13. 10.3389/fmicb.2022.1036964

Rebecchi L, Altiero T, Guidetti R. Anhydrobiosis: the extreme limit of desiccation tolerance. ISJ-Invertebr Surviv J. 2007;4(2):65–81.

Rice P, Longden I, Bleasby A. EMBOSS: The European Molecular Biology Open Software Suite. Trends in Genetics. 2000;16(6):276–7. 10.1016/S0168-9525(00)02024-2

Robert X, Gouet P. Deciphering key features in protein structures with the new ENDscript server. Nucleic Acids Res. 2014;42(W1):W320–4. 10.1093/nar/gku316

Seki K, Toyoshima M. Preserving tardigrades under pressure. Nature. 1998;395(6705):853–4. 10.1038/27576

Sievers F, Higgins DG. Clustal Omega. Current Protocols in Bioinformatics. 2014;48(1):3.13.1-3.13.16. 10.1002/0471250953.bi0313s48

Sjursen H, Bayley M, Holmstrup M. Enhanced drought tolerance of a soil-dwelling springtail by pre-acclimation to a mild drought stress. Journal of Insect Physiology. 2001;47(9):1021–7. 10.1016/S0022-1910(01)00078-6

Tamura K, Stecher G, Kumar S. MEGA11: Molecular Evolutionary Genetics Analysis Version 11. Mol Biol Evol. 2021;38(7):3022–7. 10.1093/molbev/msab120

Tanaka S, Tanaka J, Miwa Y, Horikawa DD, Katayama T, Arakawa K, et al. Novel Mitochondria-Targeted Heat-Soluble Proteins Identified in the Anhydrobiotic Tardigrade Improve Osmotic Tolerance of Human Cells. PLOS ONE. 2015;10(2):e0118272. 10.1371/journal.pone.0118272

Tang B-L, Yang J, Chen X-L, Wang P, Zhao H-L, Su H-N, et al. A predator-prey interaction between a marine Pseudoalteromonas sp. and Gram-positive bacteria. Nat Commun. 2020;11(1):285. 10.1038/s41467-019-14133-x

Wełnicz W, Grohme MA, Kaczmarek Ł, Schill RO, Frohme M. Anhydrobiosis in tardigrades—The last decade. Journal of Insect Physiology. 2011;57(5):577–83. 10.1016/j.jinsphys.2011.03.019

Yabe S, Aiba Y, Sakai Y, Hazaka M, Yokota A. Thermosporothrix hazakensis gen. nov., sp. nov., isolated from compost, description of Thermosporotrichaceae fam. nov. within the class Ktedonobacteria Cavaletti et al. 2007 and emended description of the class Ktedonobacteria. International Journal of Systematic and Evolutionary Microbiology. 2010;60(8):1794–801. 10.1099/ijs.0.018069-0

Yamaguchi A, Tanaka S, Yamaguchi S, Kuwahara H, Takamura C, Imajoh-Ohmi S, et al. Two Novel Heat-Soluble Protein Families Abundantly Expressed in an Anhydrobiotic Tardigrade. PLOS ONE. 2012;7(8):e44209. 10.1371/journal.pone.0044209

Yoshida Y, Koutsovoulos G, Laetsch DR, Stevens L, Kumar S, Horikawa DD, et al. Comparative genomics of the tardigrades Hypsibius dujardini and Ramazzottius varieornatus. PLOS Biology. 2017;15(7):e2002266. 10.1371/journal.pbio.2002266

Yu T, Cui H, Li JC, Luo Y, Jiang G, Zhao H. Enzyme function prediction using contrastive learning. Science. 2023;379(6639):1358–63. 10.1126/science.adf2465

Zhao H-L, Chen X-L, Xie B-B, Zhou M-Y, Gao X, Zhang X-Y, et al. Elastolytic Mechanism of a Novel M23 Metalloprotease Pseudoalterin from Deep-sea Pseudoalteromonas sp. CF6-2. Journal of Biological Chemistry. 2012;287(47):39710–20. 10.1074/jbc.M112.405076

Zhou Q, Xiao Q, Zhang Y, Wang X, Xiao Y, Shi D. Pig liver esterases PLE1 and PLE6: heterologous expression, hydrolysis of common antibiotics and pharmacological consequences. Sci Rep. 2019;9:15564. 10.1038/s41598-019-51580-4

