## Supplement information for "A novel metalloprotease family discovered through a tardigrade"

**Table S1. Data collection of anomalous data.**

| | ***Rv*Peti_Zn** | ***Rv*Peti_Zn** | ***Rv*Peti_Mn** | ***Rv*Peti_Mn** |
|---|---|---|---|---|
| **Data collection** | | | | |
| **Diffraction source** | Spring-8 BL44XU | | | |
| **Wavelength** | 1.28 | 1.30 | 1.80 | 1.90 |
| **Resolution range** | 49.30 – 2.10 (2.16 – 2.10) | 49.28 – 2.05 (2.11 – 2.05) | 49.56 - 1.95 (2.00 - 1.95) | 49.60 – 2.06 (2.12 – 2.06) |
| **Space group** | $P\ 3_2\ 2\ 1$ | $P\ 3_2\ 2\ 1$ | $P\ 3_2\ 2\ 1$ | $P\ 3_2\ 2\ 1$ |
| ***a, b, c* (Å)** | 72.04, 72.04, 80.46 | 72.08, 72.08, 80.30 | 72.19, 72.19, 81.33 | 72.28, 72.28, 81.31 |
| ***α, β, γ* (°)** | 90, 90, 120 | 90, 90, 120 | 90, 90, 120 | 90, 90, 120 |
| **Total reflections** | 142652 | 151563 | 311123 | 267066 |
| **Unique reflections** | 14544 (1165) | 15588 (1182) | 18106 (1103) | 15426 (1042) |
| **Redundancy** | 9.8 (9.3) | 9.7 (9.8) | 17.2 (7.2) | 17.3 (7.5) |
| **Completeness (%)** | 100.00 (100.00) | 100.00 (100.00) | 98.8 (89.3) | 98.7 (88.9) |
| $I/\sigma(I)$ | 7.6 (0.6) | 12.2 (1.3) | 24.3 (1.4) | 24.1 (1.2) |
| $R_{merge}$ **(all I+ & I-)** | 0.183 (2.942) | 0.112 (1.657) | 0.076 (1.260) | 0.066 (1.385) |
| $R_{meas}$ **(all I+ & I-)** | 0.193 (3.115) | 0.118 (1.749) | 0.068 (1.358) | 0.068 (1.485) |
| $R_{pim}$ **(all I+ & I-)** | 0.061 (1.015) | 0.038 (0.555) | 0.016 (0.489) | 0.016 (0.519) |
| $CC_{1/2}$ | 0.997 (0.560) | 0.999 (0.627) | 1.000 (0.506) | 0.999 (0.533) |

**Table S2. Primers used in this research (from 5' to 3').**

| | |
|---|---|
| Primers to linearize pET28a for *Rv*Peti | |
| Forward | ATCGTCATTTCATAACATATGGCTAGCATGACTGGTGGAC |
| Reverse | GTGGGTTAGTCGAGACTGAAAATACAGGTTTTCGTGATGATGATGATGATGGCTGCT |
| Primers to make *Rv*Peti H69A | |
| Forward | GGTGCGGCCAGCGGAACCAAGAGCC |
| Reverse | GGTCTCGGTACCTCCCGTGATGATG |
| Primers to make *Rv*Peti H76A | |
| Forward | AGCGCGTGGAACGGTTACAAGCTGG |
| Reverse | CTTGGTTCCGCTGGCATGACCGGTC |
| Primers to make *Rv*Peti D83A | |
| Forward | CTGGCGATCGCCATGAACGCCGGAG |
| Reverse | CTTGTAACCGTTCCAGTGGCTCTTG |
| Primers to make *Rv*Peti E122A | |
| Forward | AACGCGGGAAACCACTGGGATATCG |
| Reverse | AGTCCAGATGTCGGATCCATCTTGG |
| Primers to make *Rv*Peti H125A | |
| Forward | AACGCGTGGGATATCGTCATTTCAT |
| Reverse | TCCCTCGTTAGTCCAGATGTCGGAT |
| Primers to make *Rv*Peti D127A | |
| Forward | TGGGCGATCGTCATTTCATAACATA |
| Reverse | GTGGTTTCCCTCGTTAGTCCAGATG |

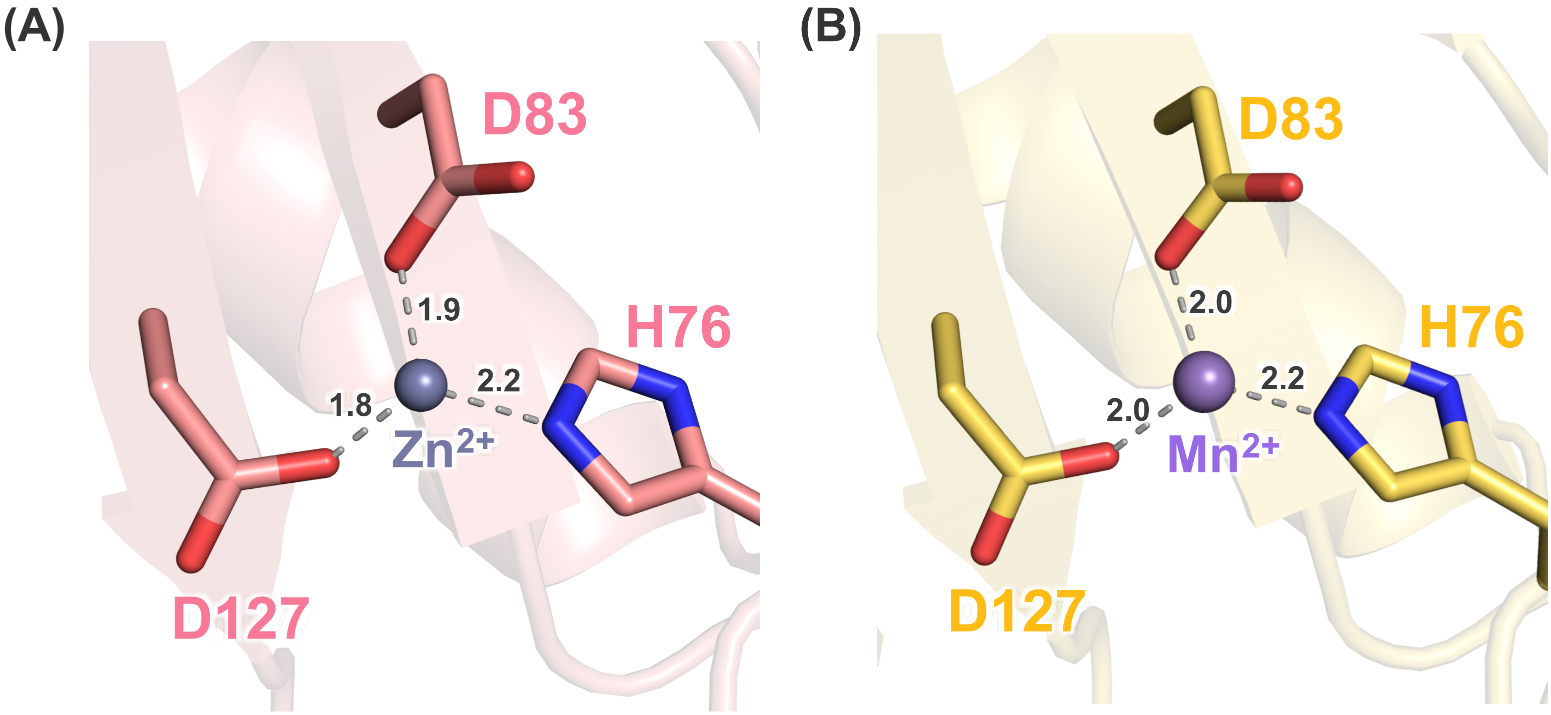


**Fig. S1 Close-up view of (A) *Rv*Peti-Zn and (B) *Rv*Peti-Mn**

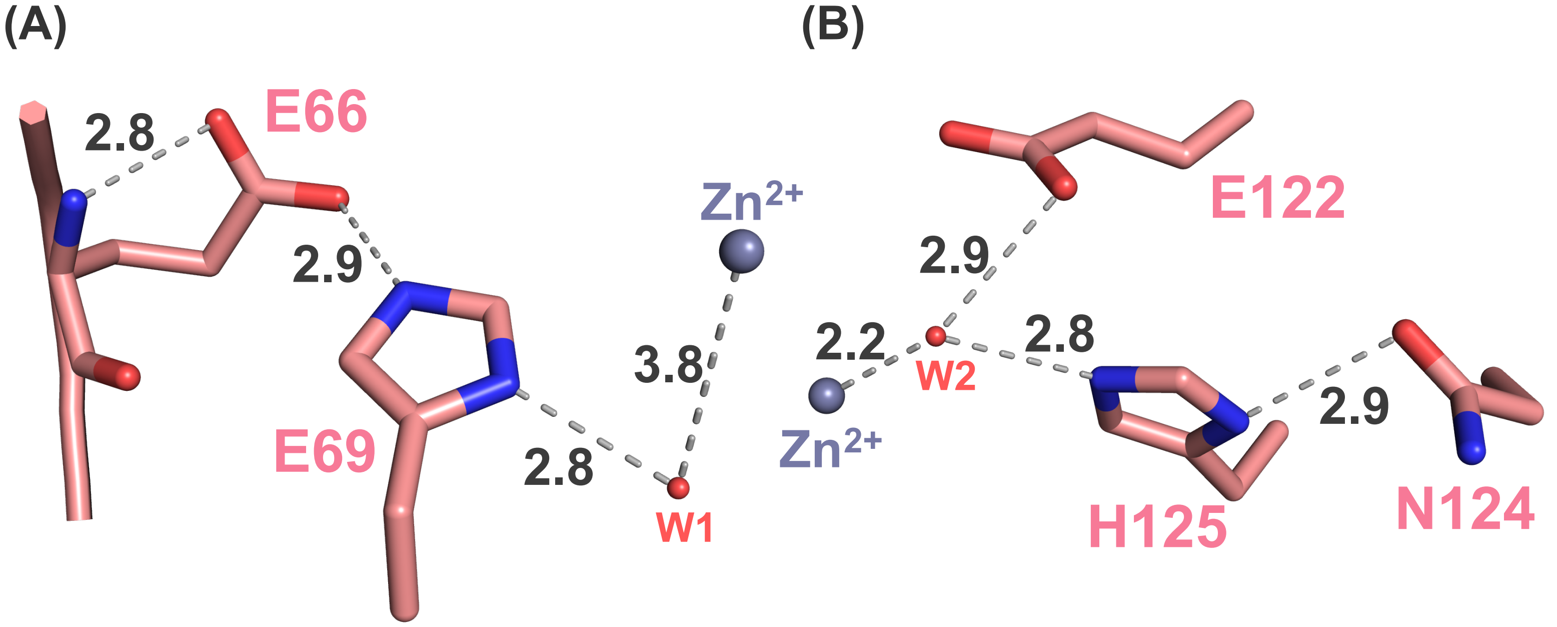


**Fig. S2 Detailed hydrogen-bond network surrounding the $Zn^{2+}$-binding site in *Rv*Peti-Zn.**

Close-up view of the metal-coordinated water molecules (W1 and W2) and their surrounding hydrogen-bond network. (A) W1 forms a hydrogen bond with His69, which in turn interacts with Glu66 and the main-chain amide nitrogen of Glu66. (B) W2 coordinates directly to the metal ion and forms hydrogen bonds with Glu122 and His125; His125 additionally interacts with Asn124.

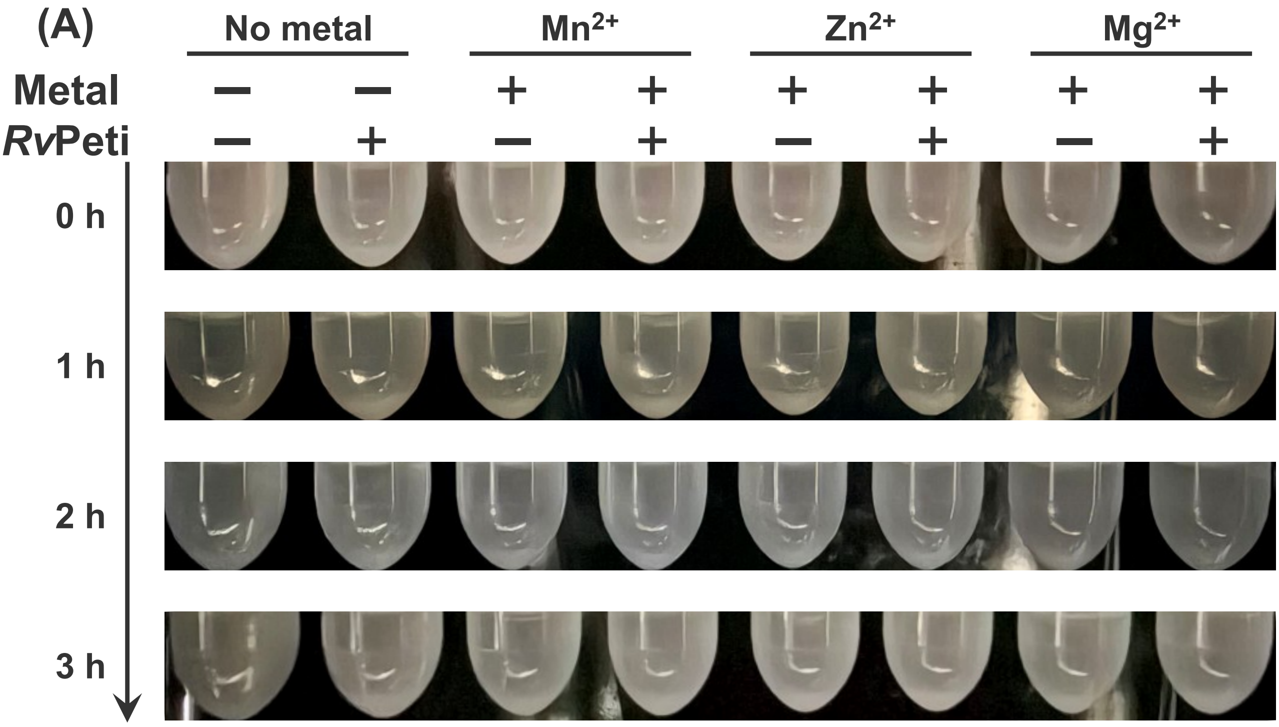


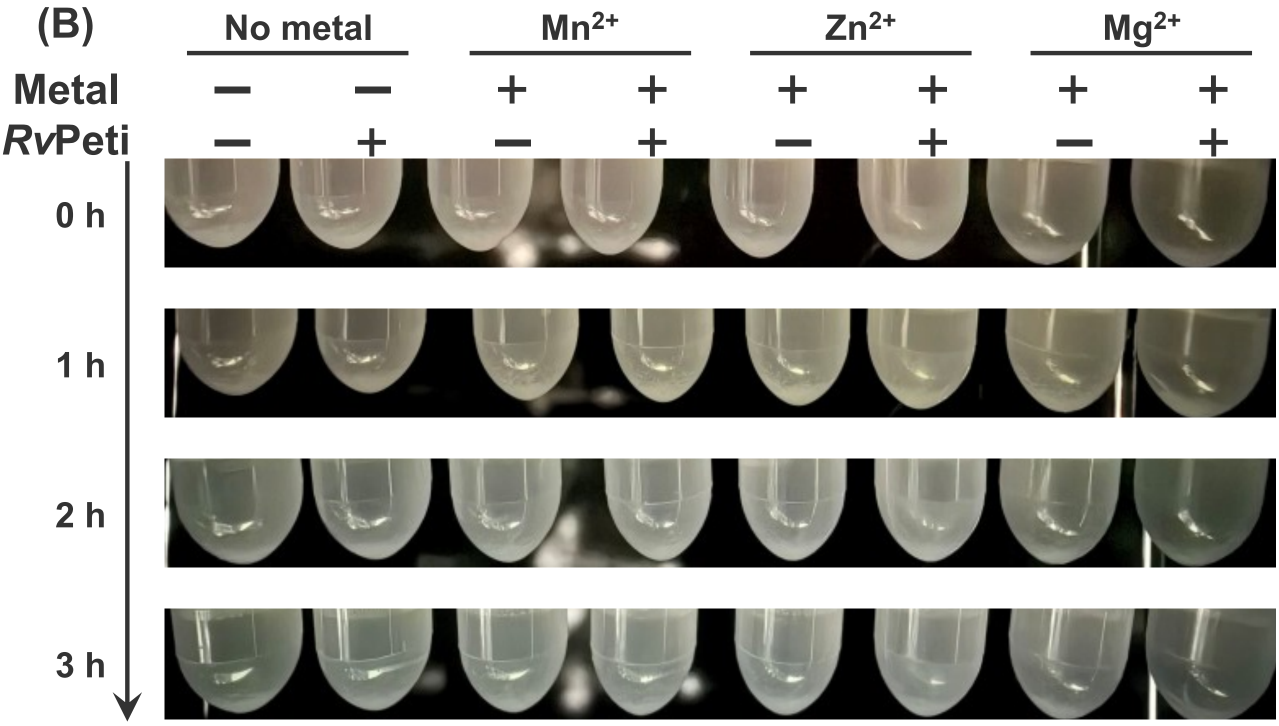


**Fig. S3** ***Rv*****Peti does not hydrolyze elastin or collagen.**
Degradation of elastin (A) and collagen (B) by *Rv*Peti. No visible degradation of either substrate was observed during incubation.

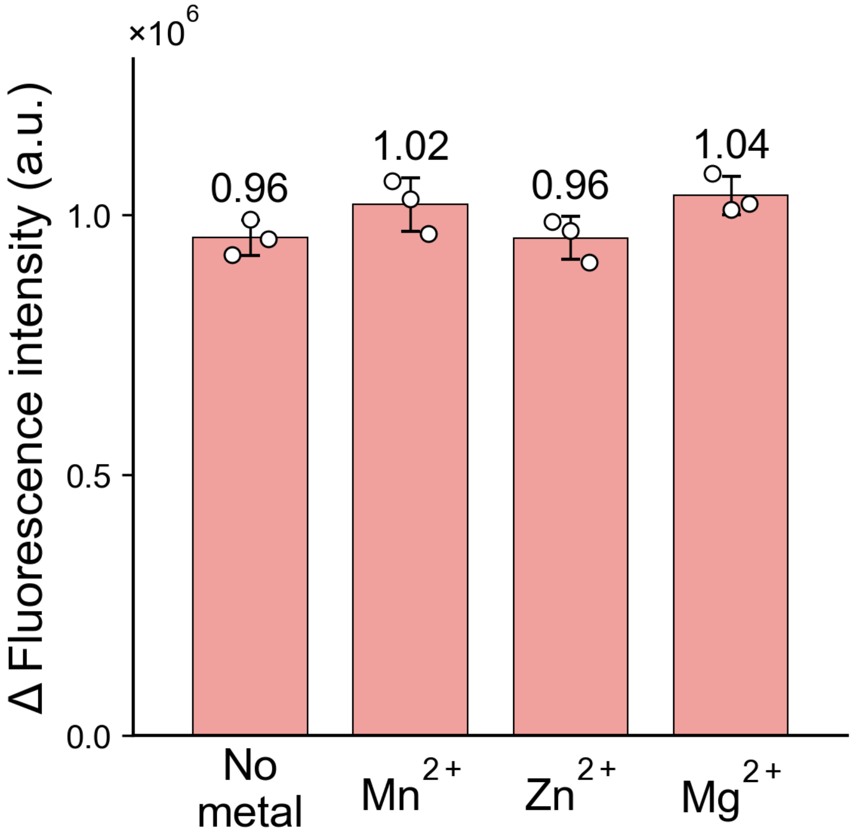


**Fig. S4 OPA assay for MDP cleavage by *Rv*Peti.**
Free amino groups generated following incubation with MDP were detected by OPA assay.

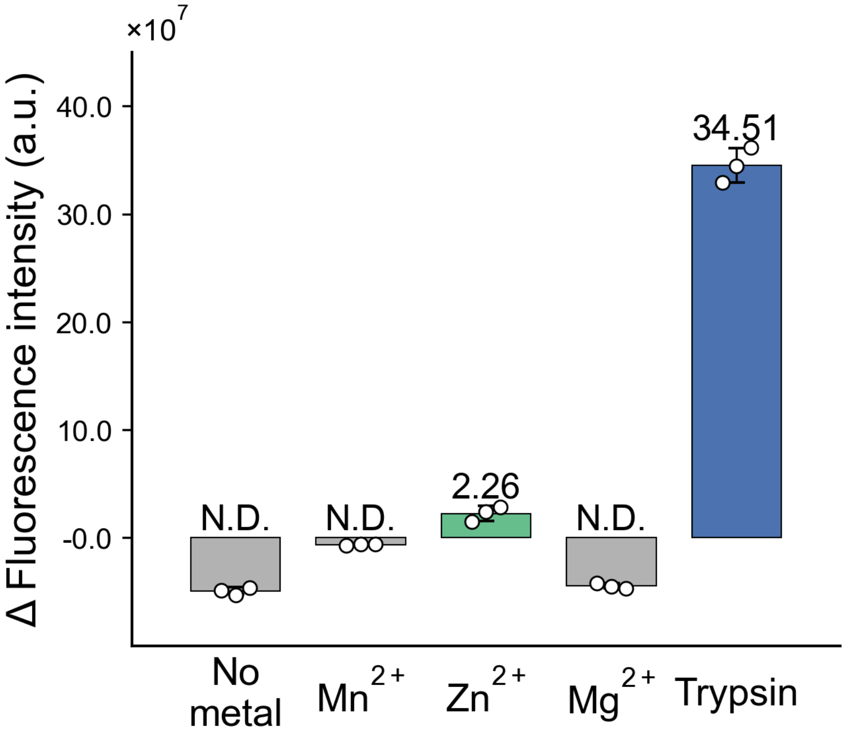


**Fig. S5 Comparison of Zn $^{2+}$-dependent proteolytic activity of *Rv*Peti with trypsin.**

Proteolytic activity toward FITC -casein was measured for *Rv*Peti in the absence of metal ions (No metal) or in the presence of $Mn^{2+}$, $Zn^{2+}$, or $Mg^{2+}$, alongside trypsin as a positive control, as in Fig. 5A.

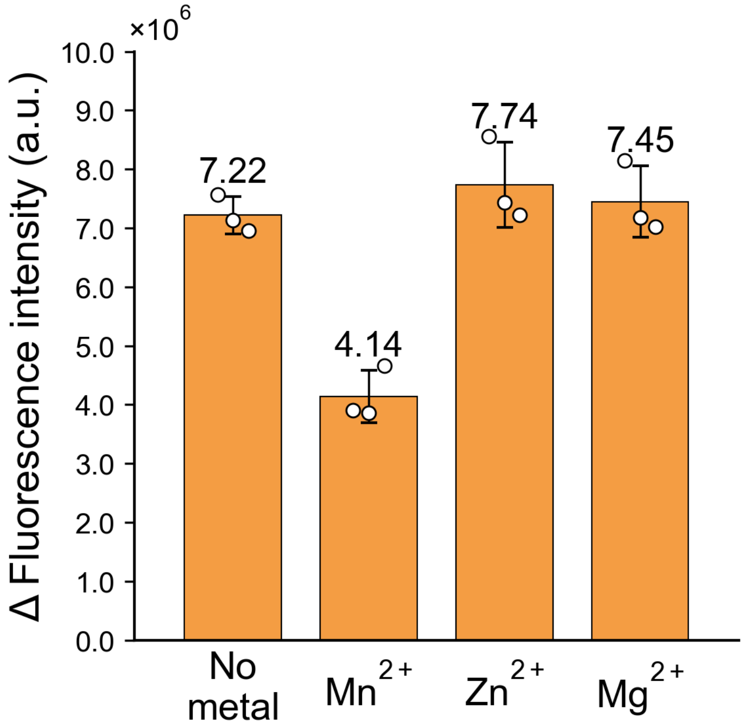


**Fig. S6 Peptidase activity of *Rv*Peti toward Gly-Gly-Gly (3Gly).**
Peptidase activity of *Rv*Peti toward 3Gly was measured in the absence of metal ions (No metal) or in the presence of $Zn^{2+}$, $Mg^{2+}$, or $Mn^{2+}$. Free amino groups generated by cleavage were detected using an OPA assay in the presence of a reducing agent. No significant difference in fluorescence was observed under $Zn^{2+}$ condition.

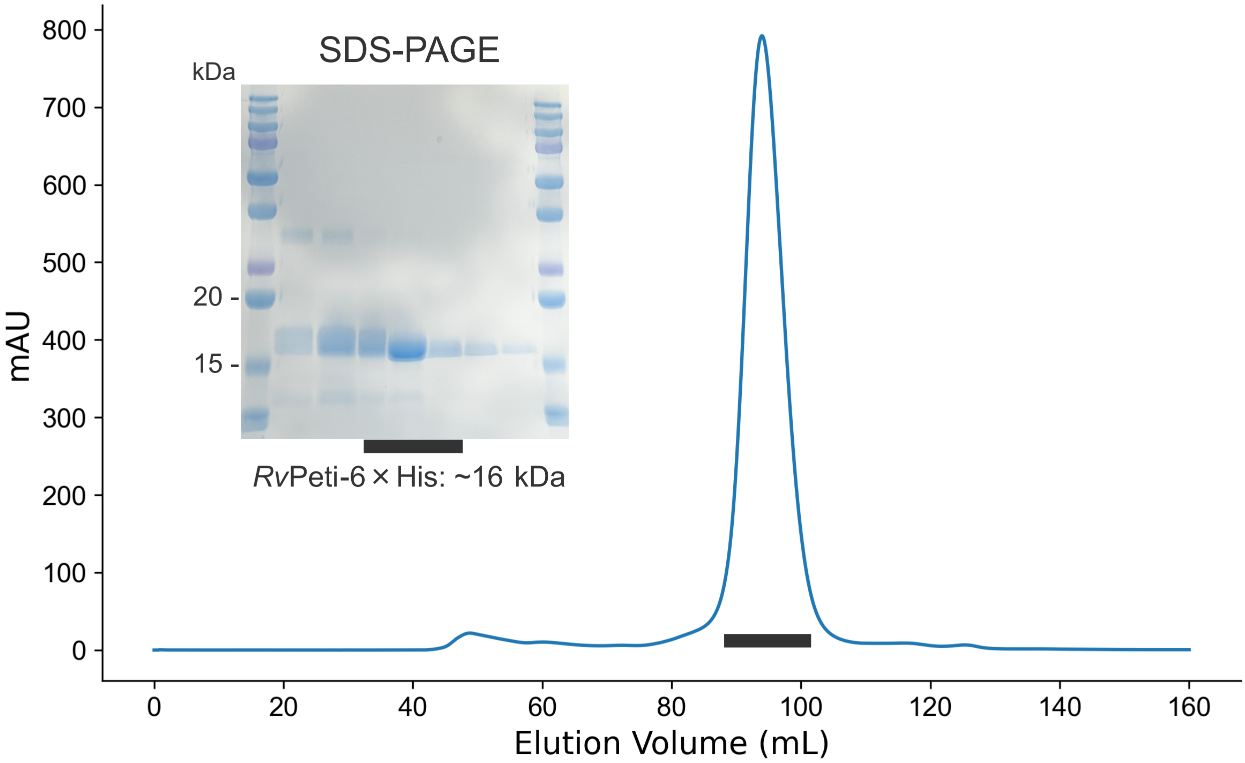


**Fig. S7 Results of size exclusion chromatography and SDS-PAGE of *Rv*Peti.**
