## Supplement Data for "A novel metalloprotease family discovered through a tardigrade"

>BAQ94592.1:1-131
tardigrade-unique protein with predicted mitochondria targeting peptide 5 [Ramazzottius varieornatus]
MSRLTHAQALQRVNAVGISVHSSGNCSDRNRPNCTSLDTILAKTIDQIVAFKRKSGKVVIITGGTETGHASG
TKSHWNGYKLDIAMNAGVTDYITKNFRYIGTTNGGRWAQYQDGSDIWTNEGNHWDIVIS

>OQV25264.1:12-139
hypothetical protein BV898_00948 [Hypsibius exemplaris]
KLSHDAAKRLLTAAGITTRSSGNCDDRENPRCTSLDQIRSETIDAVINFKRQSGARVVVTGGTEKGHAAGT
YSHWNGYKLDIAKDATADAYITGNFVYLGQLNNGWKQYKDRSDNVWTNEGNHWDVVV

>WP_111325678.1:174-300
peptidoglycan-binding domain-containing protein [Thermosporothrix hazakensis]
TKLTHSEAMSMLRANGISVVSSGNCSDRNRSNCTSLEQVRRNTIEQIIAFKQKSGCAITITGGTETGHASGT
YSHWNGYKLDIQPTSCVSNYITGHFKAAGTRGDGAKLYTDGADRIYARESNHWDI

>ODM95324.1:18-145
hypothetical protein Ocin01_11354 [Orchesella cincta]
SAVTHAEALRRFQAVGVTVTSSGGCSDRNKPTCTSFDSARENTVAGLVTLKTASGCVINVTGGTETGHASG
TYSHWNGYKADISLNTCINNYVTGHFAYIGVRGDGAPMYQSAAGNIYAKEGNHWDIL

>KAN0705896.1:29-155
hypothetical protein BAU_011521 [Basidioascus undulatus]
LTQAQAESRVRAAGILISSSGGCTDRTRPTCTSLQGVLSGTIDDIITLKKSSGCSITITGGTETGHASGAKSHS
TGYKLDISLNSCINSYIQNSFTYIGQRSDGAAQYRAASGNVYAKEGNHWDILF

>WP_246470422.1:17-145
hypothetical protein [Streptomyces olivoverticillatus]
VAKLTQSQAEARLRAAGISWTSSGHCTNRNHSNCTSFEQINVSTVNGIITFKRASGCSITITGGTETGHASGS
KSHWNGYKVDVNPTACVTDYIKRNFKYIGQRGDGAAQYQSSAGNIYARESTHWDIT

>WP_069817073.1:48-171
hypothetical protein [Streptomyces sp. TP-A0874]
AKLSHSQAMSQLNAAGIGVTSSGGCTDRNKPNCTSLDQVNSETIAGVITLKSASGCAITITGGTETGHADG
TYSHWNGYKLDLSMTSCLTNYITSAFTSIGGSKWQSAAGNIYYNEVNHWDVTF

>XP_021960849.1:18-145
uncharacterized protein LOC110856663 [Folsomia candida]
SAMTHAQAMTRLTAAGMSVTSSGGCSDRNNPTCTSFEGVRENTIAGLETLKRASACPINVTGGTETGHAS

GTYSHWNGYKADISLNACINSYVTNSFTYIGDRGDGAPMYQSAAGNIYAKEGNHWDIL

>KAL2261059.1:19-144
hypothetical protein VTK26DRAFT_4747 [Humicola hyalothermophila]
AKLTHSQATKRLRAAGIKWRSSGGCSNRNVPTCTSFDQVNKATIDAVITLKKACKCPITITGGTETGHAGG
KYSHWNGYKLDLAKNAALNRYITKNFRRITNRGDGSAQYKARSGNIYANEGNHWD

>HEY6739884.1:37-164
hypothetical protein [Actinopolymorphaceae bacterium]
VTKLSDADARARVTAAGIAVWSSGGCSNRNRPNCTSLDQVNLTTIQGILTLKSASGCAITITGGTETGHAGG
TYSHWNGYKLDISLSSCIDGYITRSFTFIGYRGDGAALYESASGNIYAREGNHWDI

>WP_356185136.1:41-170
hypothetical protein [Streptomyces sp. NPDC004647]
VTKLTHAQATSRLSAAGISWTSSGGCSDRNNPTCTSFEQVNLATIQGAETLRSASGCALTITGGTETGHASG
TYSHWNGYKLDFSPTSCVTNYVQNTFTYIGTRPGDGAPQYRSGSGNIYARESNHWDVL

>WP_035906203.1:33-161
hypothetical protein [Knoellia subterranea]
VTKLTHAQATAKLSAAGISWTSSGGCSNRNVSTCTSFDQVNSATIDGIITLKRASGCSILITGGTETGHASGT
YSHWNGYKLDISPYTCISNYITGTFPYIGLRGDGARQYKSGAGNIYARESNHWDIL

>WP_357850996.1:36-161
hypothetical protein [Micromonospora sp. NPDC006766]
VSHASATSQLSAAGISWTSSGGCSDRNRSNCTSFEGIRQATIDGIITFKRASGCAITITGGTETGHASGTYSH
WNGYKVDISKTTCVANWITSKYTYMGYISGWGYQYKAPSGNLYTDEGNHWDIL

>HEY8373693.1:34-160
hypothetical protein [Pseudonocardiaceae bacterium]
KMTHAQATARLSAAGITWSSSGGCSDRNKPNCTSFEQINSGTIDGIITLKQASGCAINITGGTETGHASGTYS
HWNGYKVDISLNSCINSYIRNSFTRIVPPSFGSEQYRSAAGNIYTLESNHWDIL

>WP_088739919.1:48-174
hypothetical protein [Haliangium sp. UPWRP_2]
AKLSQSDAAAMIRAAGISVYSSGGCTNRNRSNCTSFEQINSATVNGIITFKKASGCAVTITGGTETGHASGT
YSHWNGYKVDVSPTSCVTNYIHKNFSYIGLRGDGAPLYKSAAGNLYADEGSHWDI

>WP_420190851.1:32-160

hypothetical protein [Knoellia tibetensis]
VTKLTHSQATAKLSAAGITWSSSGGCSNRNISTCTSFDQINSTTIDGVITLKRSSGCAVNITGGTETGHASGT
YSHWNGYKVDISLYSCVSSYITNTFTYAGTRSDGAALYRSGAGNLYAKEGSHWDIL

>WP_414094395.1:27-148
hypothetical protein [Lentzea alba]
TKLTHAQATSQLRAAGITWSSSGGCSDRNRPNCTSFEQVRQTSISGAITFKRASGCALNITGGTETGHASGT
YSHWNGYKLDYSLSTCVSNYITRTFTSIGGSKWRSGSGNIYYRESNHWDV

>WP_181766814.1:41-165
hypothetical protein [Streptomyces albidus (ex Kaewkla and Franco 2022)]
LAKLSHSQAASQLSGAGISITSSGGCSDRNKPNCTSLDQVNSATIQGAITLKGASGCALTVTGGTETGHADG
TYSHWNGYKLDFHMTSCVTNYVKNTFTSIGGNKWQSGSGNIYFDEGNHWDVTF

>WP_275032578.1:36-162
hypothetical protein [Micromonospora cathayae]
ISHSSAASQLRAAGISWTSSGNCSDRNNPTCTSFEGVRQATIDGIITFKRASGCAITITAGTETGHSGGTYSH
WNGYKLDIALSTCVQNYVSTYYTYVGYISGFGYQYRAPSGNLYTKEGSHWDILF

>KAG8766804.1:26-146
hypothetical protein FRC12_006651 [Ceratobasidium sp. 428]
TRLSHAQALRILNAAGIGVTSSGHCSARNRPTCTSLDTIRCETLTGLIAFKQASRCAITVTGGTETGHENGPH
SHWNGYKVDCHMTPCLTSYIHAHFRKTDNTHWMSGRNIYHYEGNHWDI

>WP_400397715.1:38-162
hypothetical protein [Micromonospora sp. NPDC092111]
HGSAASQLSAAGIGWVSSGNCSDRNNPTCTSFEGIRQSTIDGIITFKRASGCAITITGGTETGHAGGTYSHW
NGYKLDIAKTTCVYNYISRAYTYLGYISGWGYQWRAASGNLYTDEGNHWDILF

>KAM7203318.1:19-147
hypothetical protein V8F33_002309 [Rhypophila sp. PSN 637]
AKLTHSQATSRLSAAGISWTSSGGCSNRNVATCTSFDQVREETIAGAITLKQACGCAITITGGTETGHASGT
YSHWNGYKLDMSKTTGLNNYITSTFTRIADRSDGYAQYKAGSGNIYCNEGNHWDITF

>WP_393651457.1:36-165
hypothetical protein [Streptomyces sp. NPDC048639]
VAKLTHAQATSRLRAAGISWTSSGGCSDRNNPTCTSFDQVNLATIQGAETLRSAGGCALTITGGTETGHAS
GTYSHWNGYKLDFSPTSCLSNYITGTFTNIGPRPGDGAAQYQSGSGNIYARESNHWDVL

>WP_451762057.1:35-162
hypothetical protein [Streptomyces tricolor]
TKLTHSQAAAKLRAAGVSWTSSGHCSDRNRKNCTSFSQINSGTVDGVIAFNRASGCRITITGGTETGHARG
TYSHWNGYKVDITPTTCVTRYIKRHYDYIGRRGDGAPMYRTSAGNVYADEGNHWDIT

>WP_405096288.1:36-162
hypothetical protein [Micromonospora sp. NBC_01412]
VTHSSATSQLAAAGISWTSSGGCSDRNNPTCTSFDGVRQATIEGIITFKRASGCAITVTAGTETGHIGGTYSH
WNGYKDIALSTCVQNYISAAYTYVGYISGFGYQYRAGSGNLYTKEGSHWDILF

>MIE4193390.1:36-166
hypothetical protein [Candidatus Streptomyces philanthi]
TKLTHNQAAAKLRAAGINWTSSGNCSDRNRKNCTSFTSINSGTIDGVIAFKRASGCTLTITGGTETGHASGT
YSHWNGFKVDISPATAKSCPTLYITHHYTYIGKRSDGAPMYRTSSGNTYANEGNHWDIT

>MFO0573685.1:48-174
hypothetical protein [Polyangia bacterium]
AKLSQSQAESRIRAAGISVYSSGGCTNRNKSNCTSFEQINDSTVSGIITLKSASGCAITITGGTETGHASGTYS
HWNGYKVDISITSCISSYITRNFTYQGKRGDGAPLYKSAAGNLYANEGSHWDI

>WP_364207365.1:35-160
MULTISPECIES: hypothetical protein [Polymorphospora]
ITHSSATSQLSAAGISWTSSGGCSNRNVPTCTSFDGIRQATITGIITFKRASGCAVTITAGTETGHASGTYSHW
NGYKVDIRLSTCVENYITRSYTYVGYISGWGHQYRAASGNLYTKEGSHWDIL

>WP_091648777.1:36-161
hypothetical protein [Micromonospora pallida]
ISHSSATTQLRNAGISWTSSGGCSDRNNPTCTSFDGIRQATIDGIITFKRASGCGIIITAGTETGHAGGTYSHW
NGYKVDIGLSTCIQNYISANYTYVGYISGFGYQYRAPSGNLYTKEGSHWDIL

>WP_436658524.1:36-161
hypothetical protein [Micromonospora sp. URMC 103]
ISHSSATSQLRAAGISWTSSGGCSDRNVPTCTSFDGIRQATIDGIITFKRASGCAITITAGTEVGHASGTYSHW
NGYKLDIAPSTCIQNYISSHYTYLGYISGWGYQYRAASGNLYTLEGNHWDIL

>WP_210987630.1:34-163
hypothetical protein [Streptomyces sp. F63]

VTKLTHGQATARLSAAGISWRSSGNCSDRNNPTCTSFEQVNLPTIQGAETLKSASGCALTITGGTETGHASG
TYSHWNGYKLDFSPTSCLSGYITGTFTYIGERGDGAAQYKAASGNIYARESNHWDVTF

>WP_238153931.1:16-145
hypothetical protein [Streptomyces xinghaiensis]
VTKLTHAQATARLSAAGISWRSSGNCSDRNNPTCTSFEQVNLATIQGAETLKSASGCALTITGGTETGHASG
TYSHWNGYKLDFNPTTCLSGYITGTFTYIGERADGAAQYKAASGNIYARESNHWDVTF

>WP_392845405.1:35-162
hypothetical protein [Streptomyces calvus]
TKLTHNQAAAKLRSAGITWSSSGNCSDRNRRTCTSFTRINSGTVDGIITFKRASGCKVNITGGTETGHAGGT
YSHWNGYKVDITPTTCVNRYIKREFTYIGKRGDGAAMYRSSAGNVYANEGNHWDIT

>KAK3316010.1:20-147
hypothetical protein B0H66DRAFT_275154 [Apodospora peruviana]
KLTHAQATARLSAAGISWSSSGGCSNRNVATCTSFDQVREETIAGAITLKSACGCAITITGGTETGHASGTYS
HWNGYKLDMSKTTGLNNYITSTFTRIADRSDGYAQYQARSGNIYCNEGNHWDITF

>KAB8141844.1:34-161
hypothetical protein F8S13_17045 [Chloroflexia bacterium SDU3-3]
TKLTHAQATAKLSAAGITWSSSGGCSNRNNATCTSFEQINSGTIDSIIVFKNASGCAVNITGGTETGHASGTY
SHWNGYKVDISISTCVNNWITSAYTYIGVRSDGAAMYQSAAGNIYAREGSHWDIL

>WP_291419705.1:36-156
hypothetical protein [Actinophytocola sp.]
LTHRQATAKLRAAKISWTSSGGCSNRNRPNCTSFTGIKSGTVSGIITFKKSSRCKVTVTGGTETGHASGTYS
HWNGYKVDIAKTSCTTNYIKRSFRSIGGGRWRSSAGNVYYDEGNHWDIT

>WP_375782002.1:56-184
hypothetical protein [Streptomyces zingiberis]
VTKLTHSQATTRLRNAGVSWTSSGGCSNRNQSNCTSFEQVNLATIQGAETLKSASGCALTITGGTETGHAS
GTYSHWNGYKLDYSLSTCVNNYVTGNFTYIGTRSDGAAQYRSGAGNIYAKEGNHWDVL

>WP_380088416.1:32-160
hypothetical protein [Knoellia altitudinis]
VTKLTHSQATAKLSAAGISWTSSGGCSNRNVSTCTSFEQVNSATIDGIITFKRASGCAITITGGTETGHASGT
YSHWNGYKVDIGLTTCAGNYITGTFPYNGTRGDGAPTYKSGAGNIYAKESNHWDIL

>WP_425892456.1:36-162

hypothetical protein [Micromonospora sp. DT4]

ISHSSATSQLRAAGISWTSSGNCSDRNVSTCTSFEGVRQATIDGIITFKRASGCAITITAGTEVGHAGGTYSH
WNGYKLDIALSTCIQNYISANYTYVGYISGFGYQYRAPSGNLYTKEGSHWDILF

>WP_046563892.1:36-161

hypothetical protein [Micromonospora sp. HK10]

ISHSSATSQLRAAGISWTSSGNCSDRNVSTCTSFDGIRQATIDGIITFKRASGCAITITAGTEVGHASGTYSHW
NGYKLDIALSTCIQNYVSAYYAYVGYISGFGYQYRAPSGNLYTKEGSHWDIL

>WP_388327119.1:36-162

hypothetical protein [Streptomyces cinerochromogenes]

KLTHQQAAAKLRSAGVSWTSSGHCSDRNRKTCTSFSRINSGTVDGIIAFKRASGCRITITGGTETGHAGGR
YSHWNGYKVDITPTTCVTRYIKRHYDYIGRRGDGAPMYRTPAGNVYANEGNHWDIT

>WP_135342478.1:40-168

hypothetical protein [Streptomyces palmae]

VTKITHAQATSMFREVGITWSSSGNCSDRNNSTCTSFDQLNLATAQGAQTLKRATGCALNITGGTETGHAS
GTYSHWNGYKLDYALNSCVSNYITGTFSYIGVRSDGAKMYQSGSGNIYALEGNHWDVL

>WP_344104158.1:23-148

hypothetical protein [Myceligenerans crystallogenes]

LTHAQATAQLSAAGITWSSSGNCSNRYNSTCTSFEQIRQTTIDGIKTLKRSSGCAVNITGGTETGHSSGTYSH
WNGYKVDISKYTCVGNYIKNNFTYVGYVSGWGYQYRSGAGNLYTDEGNHWDIL

>CAG7719990.1:19-145

unnamed protein product [Allacma fusca]

AKLTHSQALSQIQAAGVGISSSGGCSDRSKPTCTSLEQINVITINGVVTLKKSSGCSLTVTGGTEVGHASGT
YSHYNGYKVDYSLNTCISGYITRSFTRIADRSDGAAQYRSAAGNIYALEGNHWDV

>MEU2722090.1:20-143

hypothetical protein [Streptomyces smyrnaeus]

LAKLTHAQATAKFRAAGLTWTSSGGCSDRNNPTCTSFDGLRSATADGAITLKRASGCAVRLTGGTETGHAS
GTYSHWNGYKLDMSLSSCLTNYITGTFTSIGGSKWKSASGNIYYRESNHWDVT

>WP_392151346.1:67-193

peptidoglycan-binding domain-containing protein [Streptomyces sp. NPDC015661]

LTHSQAAAIFTAAGITWSSSGNCSDRNSSSCTSFDGLRQASADGAVALRRASGCAMNITGGTETGHASGTY

SHWNGYKLDFAMSSCLTGYITGTFPYSGQRGDGAALYTAPSGNIYANEGNHWDVTF

>WP_281575206.1:36-161
hypothetical protein [Micromonospora sp. WMMD812]
ITHSSATSQLRAANISWTSSGNCSDRNNPTCTSFAGIRQATIDGIITFKRASGCAVTITAGTETGHAGGTYSH
WNGYKLDIALSTCIQNYISSHYTYLGYISGWGYQYRAASGNLYTKEGSHWDIL

>WP_132820608.1:37-164
hypothetical protein [Streptomyces hainanensis]
TKITHSQATSMFSSAGVTWSSSGGCSNRNVPTCTSFDQLNLATAQGAQTLRRASGCALNITGGTETGHASG
TYSHWNGYKLDFALNSCVNSYVTNNFAFIGYRGDGAAQYQSGSGNIYAREGNHWDVT

>WP_353950580.1:32-160
hypothetical protein [Knoellia sp. S7-12]
VTKLTHSQATAKLSAAGITWSSSGNCSNRNVSTCTSFDQINSTTIDGIITLKRSSGCAINATGGTEVGHSSGT
YSHWNGYKIDISLYSCVGSYITRTFPYAGTRGDGAPMYKSGAGNIYAKESNHWDIL

>MEV8363218.1:17-141
hypothetical protein [Streptomyces niveus]
LAKLSHSTAASQLSGAGIGISSSGGCSNRNNPTCTSLDQVNSATIQGAITLKNASGCGLTVTGGTETGHASG
TYSHWNGYKLDFSMTSCLTNYVTRNFTSIGGSKWQSGAGNIYYNEVNHWDVTF

>KAK4216428.1:22-148
hypothetical protein QBC37DRAFT_385328 [Rhypophila decipiens]
KLSHSQATAKLSAAGVKWMSSGGCSDRNRPTCTSFTNVREETINGAITLKRACGCALTITGGTETGHASGT
YSHWNGYKLDFKKASGINNYITSTFTRIADRGGYPAYKAKSGNIYVNEGNHWDVTF

>WP_443072816.1:58-182
peptidoglycan-binding domain-containing protein [Streptomyces sp. WMMC897]
KLTHAQAAAMFQSAGISWSSSGNCSNRYNSSCTSFDQLRRTSADGIIAFQRASGCTVHITGGTETGHASGT
YSHWNGYKIDFAPYSCVSDYVTGTFDYLGTRSDGAALYESGAGNVYAHEGSHWD

>WP_147061574.1:33-160
hypothetical protein [Knoellia locipacati]
TKLTHAQATAQLSAAGITWSSSGGCSNRNVSTCTSFDQVNSATISGVVTLKRSSGCAINLTGGTETGHASGT
YSHWNGYKVDIALATCINNYVTNTFAYAGVRSDGATLYRSGAGNIYAKEGNHWDVL

>XP_037033138.1:25-149

uncharacterized protein LOC119072086 [Bradysia coprophila]
THAEALEKIEAVGITIASSGNCSDRNIASCTSLEQIRRKTIDSGVIVLKEASQCDIIITGGTEVGHSSSTYSHFN
GYKLDYRLNTCLNDYIMKRYTYLGLRGDGAKKYMSAAGNIYALEGNHWDV

>WP_028846457.1:45-168
MULTISPECIES: hypothetical protein [Thermocrispum]
VTKLSHAEATKMLRAAGITWSSSGNCSDRNKPNCTSFEQVNLTTIQGIQTLKRASGCAINITGGTETGHASG
TYSHWNGYKLDTSLSTCLNNYIQRSFTSIGGNKWRSASGNIYYKEGNHWDIT

>CAF0866508.1:25-151
unnamed protein product [Adineta steineri]
THAAALTQIKSVGIGISSSGGCSNRANPKCTSLDKVRCNAIRCLKTLKTSSGCALTVTGGTETGHAGGTYSH
GNGYKIDISLNACINSYITKSFAYIGKRGDGAAQYKASSGNLYAKEGNHWDITFT

>WP_376770735.1:41-163
hypothetical protein [Streptomyces physcomitrii]
FAKLSHSQAASQLSNAGIGISSSGGCSDRNTPTCTSLEQVNSATIQGAITLKGASGCSLTVTGGTEVGHASGT
YSHWNGYKLDFSMNSCLTNYITGTFTSIGGSKWQSGSGNIYYNESNHWDV

>WP_416291864.1:36-161
hypothetical protein [Micromonospora sp. SL4-19]
ITHSSATSQLSAAGIGWVSSGNCSNRNNPTCTSFEGIRQATISGIITFKRASGCAITITGGTETGHAGGTYSHW
NGYKVDIAKTTCVYNWITSAYTYVGYISGWGYQWRAPSGNLYTDEGNHWDIL

>WP_374226970.1:27-151
hypothetical protein [Streptomyces sp. JJ38]
KLTHAQAAAMFQSAGISWSSSGNCSDRYNSSCTSFDQLRRTSADGIIALKRASGCTVHITGGTETGHASGT
YSHWNGYKIDFAPYSCVSNYVTGTFTYLGKRSDGAALYESGSGNVYAHEGSHWD

>WP_091560868.1:36-161
hypothetical protein [Micromonospora pattaloongensis]
VSHSSATSQLRAAGITWSSSGGCSDRNTPTCTSFEGIRQTTIDGIITFKRASGCAVNITGGTETGHASGTYSH
WNGYKVDTSLSTCIENYIKAHYTYVGYVSGWGYQYRAPSGNLYTKEGSHWDVL

>WP_203845263.1:35-160
hypothetical protein [Dactylosporangium siamense]
VSHSSAASQLRAAGISWTSSGNCSDRNNATCTSFEGIRQATITGIITFKRASGCAVTITAGTETGHAGGTYSH
WNGYKLDIQKTSCVNGYITGHYAYVGYIPGWGYQYQAASGNLYTDEGNHWDIL

>KFH44300.1:16-145

hypothetical protein ACRE_049130 [Hapsidospora chrysogenum ATCC 11550]

QSKLSHSQAVSQLSAAGISVSSSGGCSDRNTPTCTSLEQVNADTVAGAVTLKGACGCTVRVTGGTETGHA
GGTYSHWNGYKLDFSKTSGLNSYITNTFTRIADRGDGYPQWEAASGNIYCDEGNHWDVTF

>WP_212759321.1:33-158

hypothetical protein [Promicromonospora citrea]

LTHAQATSQLSAAGISWTSSGGCSDRNVATCTSFDQVRQTTIDGIKTLRSATGCAITITGGTETGHSSGTYSH
WNGYKLDIRHSTCLDSYITSTFTYVGYISGWGHQYRSASGNLYTDEGNHWDIL

>WP_334631209.1:36-162

hypothetical protein [Micromonospora sp. CPCC 205539]

ITHASATSQLSAAGISWTSSGNCSNRNVATCTSFTDVRQATISGIITFKRASGCAITVTAGTEVGHAGGTYSH
WNGYKLDIALSTCIQNYISAHYTYIGYIAGFGYQYQAASGNLYTKEGSHWDILF

>WP_433107602.1:50-173

hypothetical protein [Streptomyces sp. CA-250714]

LAKLTHAQATAKFRAAGITWTSSGGCSDRNNPSCTSFEQLRSETADGAITLKQASGCAIRLTGGTETGHASG
TYSHWNGYKLDMSLTSCLDNYVTRTFTSIGGSKWKSPSGNVYYREGNHWDVT

>WP_051722540.1:39-168

hypothetical protein [Streptomyces albus]

VTKLTHAQATARLSAAGISWRSSGNCSDRNNPTCTSFEQVNLATIQGAETLKSASRCAVTITGGTETGHASG
TYSHWNGYKLDFNPTTCLSGYITGTFTYIGDRADGAAQYKAASGNIYARESNHWDVTF

>CAR3061427.1:33-158

Peptidoglycan-binding protein [Catelliglobosispora sp.]

LTHAQATAQLSAAGITWSSSGNCSDRNNPTCTSFTNVRQTTISGIKTLKTYSGCAINITGGTETGHSGGTYS
HWNGYKLDISKYTCVGNYIKNNFTYVGYISGWGYQWRSGAGNLYTDEGNHWDIL

>WP_406698289.1:1-130

hypothetical protein [Singulisphaera photorubra]

MAKLSHADAASKLQAAGITWSSSGNCSDRNNKRCTSFDEINSETIDGIISFKKVSGCEVRITGGTETGHSTS
TKSHWNGFKVDINPSVCVSDFIENNFEAAGVRSDGAKMFKDGAGNTYARESDHWDITF

>WP_212841993.1:33-158

hypothetical protein [Catellatospora sp. IY07-71]

LTHAEATAQLRAAGITWSSSGNCSDRNNPTCTSFTNVRQTTISGIKTLKTASGCAINITAGTETGHSSGTYSH
WNGYKLDISKYTCIGNYIKNNFTYVGYISGWGYQYRSGAGNLYTDEGNHWDIL

>WP_377234797.1:33-158
hypothetical protein [Promicromonospora sp. NPDC059942]
LTHAQATAQLSAAGISWTSSGGCSNRNVATCTSFDQVRQTTIDGIKTLRSASGCAITITGGTETGHSSGTYSH
WNGYKLDTRQSTCIDSYISNNFTYVGYISGWGYQYRSGAGNLYTDEGNHWDIL

>WP_361478370.1:37-164
hypothetical protein [Streptomyces sp. NPDC019937]
TKLTHAQAASRLRAAGISWTSSGHCSNRNNRHCTSFTQINSGTVKGITTFKQSSHCAITITGGTETGHASGR
YSHWNGYKVDIAPTSCVTNYIKRNFRYVGRRGDGAPMYKSSAGNIYARESSHWDIT

>WP_343446389.1:36-161 hypothetical protein
[Micromonospora schwarzwaldensis]
VSHSSASSQLRAAGIGWVSSGNCSDRNTPTCTSFEGIRQSTIDGIITFKRASGCGITITAGTETGHAGGTYSH
WNGYKLDIGKSTCINNYISQAYTYMGYISGWGYQWRAPSGNLYTDEGNHWDIL

>WP_203934977.1:36-163
hypothetical protein [Virgisporangium ochraceum]
RLTHTQAVKQLDAAGISWDASGDCTDRDRPACTSFSGIRQSTIDGIRTLRKASGCPLVVTGGTETGHASGTY
SHWNGWKIDIRRTACVDAYVGKWFTLVGKVEGWGTQYKAKSGNLYTNEGHHWDILF

>XVZ78898.1:36-161
hypothetical protein ACQPWV_11960 [Plantactinospora sp. CA-291049]
ISHASATSQLSAAGISWTSSGGCSDRNNPTCTSFEGIRQATITGIITFKQASGCAITITGGTETGHANGTYSHW
NGYKVDFGLTTCAQNYISAYYTYLGYVSGWGYQYQAPSGNLYTREGNHWDVL

>WP_343759297.1:47-174
hypothetical protein [Deinococcus depolymerans]
TRLSDATARSRVTAAGIPVVSSGNCSVRSNASCTSLEQIYSGTIDGIITLKQASGCAVNITGGTEVGHASGTY
SHYNGYKVDTSITTCVSNYITANFTYIGVRGDGAPQYRSAAGNIYARESNHWDIL

>WP_028850985.1:42-165
hypothetical protein [Thermocrispum municipale]
VTKLSHSDATQRLRASGITWSSSGNCSDRNNPTCTSFEQVNLTTIQGIQTLKSASGCAINVTGGTETGHASG
TYSHWNGYKLDTSLSTCLNNYIQNSFSYIGGSKWQSASGNIYYKEGNHWDIT

>HEX5598549.1:36-161

hypothetical protein [Micromonosporaceae bacterium]

ISHSSATSQLRAAGITWSSSGNCSDRYNSTCTSFEGIRQVTIDGIITFKRASGCAINITGGTEIGHASGTYSHW
NGYKVDISKNSCVYNYISRYYTYVGYISGWGYQWRAPSGNLYTDEGNHWDIL

>WP_362442209.1:61-191

hypothetical protein [Streptomyces sp. NPDC048636]

LAKLSHADATGRLRNAGISWTSSGGCSDRNQPTCTSFEQVNLATIQGAVTLKQASGCALTITGGTETGHAS
GTYSHWNGYKKLDFSPTSCLSGYITRTFTNIGPRPGDGAQQYRSGAGNIYARESNHWDVLF

>WP_415645921.1:19-141

hypothetical protein [Stackebrandtia soli]

AKLTHAQAASRLSAAGITWSSSGGCSDRNNPTCTSFTQINSATISGIITLKSASGCAINVTGGTETGHADGTY
SHWNGYKLDTSLSTCLQNYITGTFTSIGGSKWQSGSGNIYYLEGNHWDIT
